# Structures and zinc ion transport pathways of the human SLC39A family of metal transporters

**DOI:** 10.1101/2025.08.09.669477

**Authors:** Xilan Wang, Christian D. Lorenz, Christer Hogstrand, Wolfgang Maret

## Abstract

The SLC39A (ZIP) family of zinc ion transporters play a pivotal role in maintaining zinc homeostasis, which is essential for numerous physiological processes involving enzyme catalysis, protein structure and regulation in signal transduction. This investigation employed AlphaFold3 to predict and analyze the 3D structures of all 14 human ZIP family members and revealed key structural and functional features, including transmembrane domains with eight alpha-helices, extracellular and cytoplasmic domains, dimerization, and zinc ion transport pathways. Unique zinc-binding motifs—composed of histidine, aspartic acid, and glutamic acid—were identified. They facilitate zinc ion attraction, selection, and transport. The findings highlight significant structural diversity in these proteins, with additional alpha-helices, disulfide bonds, and other conserved motifs that together contribute to functional specialization across the ZIP family members. Compared to predictions, which exist only for ZIP4, the models incorporate dimeric structures, rationalize loop conformations, and achieve a higher resolution. The predicted 3D structures offer enhanced insights into zinc ion transport mechanisms and provide a foundation for future research into the structural biology of these proteins and their interacting partners in physiology and pathology.

## 1. Introduction

### 1.1 Zinc transporters

Central to maintaining cellular zinc homeostasis are the SLC39A (ZIP) and the SLC30A (ZnT) families of membrane transporters, which mediate zinc influx and efflux, thereby regulating intracellular zinc ion concentration [1]. Adequate zinc transport is vital for normal growth, development, and tissue repair [2,3]. Disruptions in ZIP transporter functions can precipitate either zinc deficiency or toxicity, with profound implications for health [2–4]. Mutations in ZIP or ZnT transporter genes are linked to a range of health disorders [5–7]. The human ZIP family, which is the subject of this article, encompasses 14 members (ZIP1-14), each of which is implicated in distinct tissue-specific cellular functions [8,9].

### 1.2 Functional motifs in ZIP transporters

ZIP (Zrt-, Irt-like Protein) transporters are integral membrane proteins [10]. They have a conserved architecture of eight transmembrane (TM) α-helices, with both N- and C-termini facing the extracellular environment [10]. Defining features of ZIP transporters are the presence of histidine-rich (HIS-rich) regions, often localized in the intracellular loop between TM3 and 4, and α-helices in extracellular loops, which have a critical role in coordinating metal ions such as zinc ions [10–12]. Notably, ZIP8 and 14 are capable of transporting both zinc and other metal ions, including iron and manganese, illustrating the functional diversity within the ZIP family [13,14].

ZIP4, a key zinc transporter in the intestinal uptake of zinc, exemplifies the main features, where the HIS-rich motif in its N-terminal region is essential for high-affinity zinc binding [12]. In ZIP transporters such as ZIP1,4, and 10, specific histidine residues within the transmembrane domain (TMD) have been identified as crucial for zinc binding, selectivity, and transport efficiency [10,12,15,16]. In addition to these regions, ZIP family members harbor several other conserved motifs, such as the HNP (Histidine-Asparagine-Proline) motif located in TM2, and the ZIP signature motif in TM4 (HXXHXDH), both of which have been implicated in metal ion binding and translocation [10,12,17]. The mammalian ZIP family has varying degrees of sequence similarity among its members and accordingly is categorized into four subfamilies based on phylogenetic analyses: subfamilies I (ZIPI), II (ZIPII), gufA, and LIV-1. Subfamily I and gufA have only one member each: ZIP9 and 11, respectively; Subfamily II has ZIP1,2, and 3. The LIV-I subfamily contains ZIP4-8-,10, and 12-14. Members within the same subfamily share higher sequence similarity, reflecting conserved structural motifs and functional characteristics [9,10]. The LIV1 subfamily contains a conserved metalloproteinase-like motif (HEXPHEXGD) located on TM5, which forms the main zinc ion transport site within the TMD [18].

Although ZIP transporters share the common function of mediating zinc uptake, they display significant structural and functional diversity across the family [10,17]. Differences in the organization of TMDs contribute to this variability, allowing individual members to adapt to distinct physiological roles [10]. Despite these advances, the understanding of ZIP transporter structures and mechanisms remains incomplete, particularly with respect to their conformational dynamics, substrate specificity, and metal ion transport pathways, as there is no directly determined eukaryotic ZIP 3D structure available for any of the members of the family.

### 1.3 The structural features in the ZIP family

Two ZIP-related 3D structures have been characterized: a bacterial Zip from *Bordetella bronchiseptica* (BpZIP) with (PDB ID: 7Z6N) and without (PDB ID: 8CZJ) bound metal ions and the extracellular domain (ECD) of *Pteropus alecto* (black fruit bat) ZIP4 (PDB ID: 4X82), which shares 68% residue identity with human ZIP4 (hZIP4). hZIP4 and BbZIP share 18% identical and 36% similar residues in the TMD. The BbZIP structure reveals eight TM α-helices forming two bundles, the transport channel contains TM2, 4, 5, and 7 while the outside helices contain TM1, 3, 6, and 8 [19]. ZIP transporters operate via an elevator-type mechanism, where a mobile transport domain made up by one four-helix bundle shifts relative to a stationary scaffold made up by another four-helix bundle to enable alternating access to metal-binding sites. Structural analyses of prokaryotic ZIPs support this model, identifying highly conserved histidine residues in TM4 and 5 as essential for metal coordination and transport [19,20]. Soaking experiments of crystals with metal ions indicated that the M1 site serves as the primary transport site, while the M2 site plays a regulatory or auxiliary role. The binuclear metal center in the transport site in BpZip is bridged by the residue GLU181 on TM4. The Cd^2+^ ion at M1 is coordinated with HIS177, GLU181, GLN207, and GLU211, while the Cd^2+^ ion at M2 is coordinated with ASN178, GLU181, ASP208, and one water molecule [17,19,21]. Recent work pointed out that BpZIP works through two main processes to release metal ions into the periplasm, namely using a combination of hinge movement and sliding within its TMD. A key part of this system is a metal site called M3 site, which helps controlling the transport process. At high zinc concentrations the M3 site is an inhibitory site stopping further transport. It stabilizes the inward-facing conformation, which slows down the release of metal ions. The zinc ion at the M3 site is coordinated by HIS149 and 151 from the second intracellular loop (IL2), as well as by ASP144 from TM3 and GLU276 from TM7 [22].

From the BbZIP crystal structure in the inward-facing conformation, two models have been proposed for the transport: (1) an alternating entry mechanism, where metal ions bind at the transport (M) sites on one side while blocking the opposite side, and (2) a transient open state where transport (M) sites are briefly accessible from both sides, resembling a channel [19]. Additionally, recent research suggests that certain ZIP members (ZIP2,3,8, and 14) facilitate the co-transport of bicarbonate (HCO₃⁻) alongside metal ions, contributing to ion homeostasis and charge balance across membranes [23–25]. Bicarbonate inhibits pufferfish *Fr*ZIP2-mediated zinc uptake without affecting the pH of the buffer or the concentration of calculated free Zn(II) ion concentrations [24]. However, zinc uptake by human ZIP2 is stimulated by the same concentrations of bicarbonate, indicating different mechanisms of zinc transport, including the possibility of a proton-driven mechanism [24,26]. ZIP8 also facilitates the cellular uptake of hydrogen selenite (HSeO ^−^) through a mechanism dependent on both zinc and bicarbonate ions, with bicarbonate likely serving as a counter-ion to balance the charge during co-transport of metal cations and hydrogen selenite; although the investigation focuses on ZIP8, it also highlights the structural and functional similarity to ZIP14, suggesting its potential involvement in bicarbonate-coupled transport [25].

### 1.4 AlphaFold

AlphaFold 1 (2018) and 2 (2020) represented a transformative advancement in computational biology and structural bioinformatics. These state-of-the-art tools enables the prediction of protein structures within hours or days, a process that traditionally spans months or even years when employing experimental techniques such as Cryo-EM (cryo-electron microscopy) and X-ray crystallography. The accelerated pace of AlphaFold-driven research facilitates rapid hypothesis testing and structural analysis, significantly expediting workflows. Furthermore, the computational demands of AlphaFold are relatively modest compared to the substantial financial and labor costs associated with experimental methods. While Cryo-EM, X-ray crystallography, and NMR spectroscopy require sophisticated equipment, expensive reagents, and considerable expertise, AlphaFold relies primarily on high-performance computing resources [27–30].

The recent opening of the resource AlphaFold3 is particularly advantageous for studying proteins or complexes that pose significant challenges for experimental techniques due to issues with sample preparation or inherent instability. Its capability to process large datasets allows for efficient prediction of numerous protein structures or entire proteomes within a condensed timeframe, which is invaluable for exploring protein structures across diverse biological contexts and for large-scale drug discovery endeavors. By leveraging extensive sequence data and evolutionary information, AlphaFold3 not only enhances our understanding of protein function and stability but also provides insights that may elude traditional experimental methods [27,29].

In this investigation, we employed AlphaFold3 to predict the structures of all 14 human ZIP family transporters and to identify the binding sites for the zinc ion. This computational approach enabled us to elucidate the unique structural features of each human ZIP transporter, highlighting both their distinctive characteristics and commonalities across the family. By providing a comprehensive analysis of these predicted structures, this work offers significant insights into the architecture and function of zinc ion transporters. Furthermore, our findings have the potential of advancing the understanding of ZIP transporters in both basic research and clinical contexts, including drug design and development of therapeutics.

Our predicted ZIP protein structures have several key advantages over those predicted by AlphaFold2 and other online protein prediction platforms. While AlphaFold2 predominantly predicts monomeric structures, which fail to reflect the dimeric configuration crucial for ZIP protein functionality [27], our AlphaFold3 models accurately represent ZIP proteins as dimers, better capturing their native conformation. Additionally, our structures explicitly incorporate zinc ion binding sites, absent in AlphaFold2 predictions, thus enhancing their utility for studying zinc transport mechanisms. Furthermore, AlphaFold2-predicted models often display instability in key regions, such as irrationally positioned loop structures, whereas our models rationalize and stabilize these conformations in simulated aqueous environments, resulting in biologically relevant structures. Our models demonstrate superior alignment and higher resolution, enabling detailed insights into zinc ion pathways and residue-specific interactions. By addressing functional considerations specific to the ZIP family, including conserved motifs critical for zinc ion binding and transport, our models capture structural variations and functional diversity across ZIP family members. These advancements make our models more accurate, detailed, and biologically relevant than AlphaFold2 predictions, filling critical gaps and advancing the understanding of ZIP protein mechanisms.

## 2. Methods

### 2.1 AlphaFold3 structural predictions

We retrieved the full-length amino acid sequences for each of the human SLC39A transporter family members from the UniProt database (https://www.uniprot.org/) and stored them in FASTA format (Table 1). To predict the 3D structures of these transporters, we utilized AlphaFold3, a state-of-the-art deep-learning-based computational tool developed by DeepMind, which achieves high accuracy in protein structure prediction by integrating sequence-based modeling with physicochemical constraints [27,30,31].

**Table 1.** Human ZIP sequences used for structural prediction.

| **Name** | **Subfamily** | **Sequence Name** | **Length (a.a.)** |
| --- | --- | --- | --- |
| ZIP12 | LIV1 | Q504Y0 | 654 |
| ZIP4 | LIV1 | Q6P5W5 | 647 |
| ZIP8 | LIV1 | Q9C0K1 | 460 |
| ZIP14 | LIV1 | Q15043 | 492 |
| ZIP10 | LIV1 | Q9ULF5 | 831 |
| ZIP6 | LIV1 | Q13433 | 755 |
| ZIP13 | LIV1 | Q96H72 | 371 |
| ZIP7 | LIV1 | Q92504 | 469 |
| ZIP5 | LIV1 | Q6ZMH5 | 540 |
| ZIP1 | ZIPII | Q9NY26 | 324 |
| ZIP2 | ZIPII | Q9NP94 | 309 |
| ZIP3 | ZIPII | Q9BRY0 | 314 |
| ZIP9 | ZIPI | Q9NUM3 | 307 |
| ZIP11 | gufA | Q8N1S5 | 342 |

To accurately model the ZIP transporters’ ability to bind divalent metal ions, particularly zinc, we incorporated eight Zn^2+^ ions into each dimeric structure, based on prior biochemical and structural evidence suggesting multiple zinc-binding sites within ZIP transporters [32]. The inclusion of zinc ions was achieved through the metal-binding site prediction capability of AlphaFold3, ensuring the structural relevance of the modeled conformations. The protein structure with the bound Zn^2+^ ions for each ZIP transporter was produced using the AlphaFold3 server (https://alphafoldserver.com/).

To further enhance structural reliability, multiple independent AlphaFold3 predictions were generated for each sequence, allowing for consensus validation of metal-coordinating residues and dimerization interfaces. Structural quality was assessed using predicted local distance difference test (pLDDT) scores and predicted aligned error (PAE) maps, ensuring confidence in both the overall fold and the accuracy of the Zn^2+^ coordination sites. A pLDDT value exceeding 90 indicates high-confidence modeling with precise structural accuracy. Scores ranging from 70 to 90 suggest reliable backbone predictions, though minor variations may exist in finer structural details [33]. The resulting structural models were visualized and analyzed using VMD (Visual Molecular Dynamics) with a focus on transmembrane topology, dimerization interfaces, and Zn^2+^ coordination geometry.

By integrating AlphaFold3’s advanced structure prediction with explicit modeling of zinc ion binding, we aimed to achieve high-resolution structural insights into all members of the human ZIP transporter family, facilitating a deeper understanding of their metal transport mechanisms and dimeric assembly.

AlphaFold3 demonstrates a strong ability to predict relatively stable structures, particularly TMDs. However, highly flexible regions, such as loops and extended N-terminal or C-terminal regions, generally exhibit lower confidence scores, often falling below 50%. This reduced confidence arises from the inherent difficulty in accurately positioning these flexible regions within the structural model [31].

To assess the reliability of the predicted structures, we present a summary of confidence scores in Table 2. The key metrics used for evaluation are defined as follows:

1. Fraction_disordered: Ranges from 0 to 1, representing the proportion of residues predicted to be disordered within the structure. A higher value indicates a greater proportion of disordered regions.
2. pTM (Predicted TM-score): Ranges from 0 to 1, reflecting the template modeling (TM) score of the predicted structure. A higher score suggests greater structural reliability.
3. has_clash (Atomic Clash Indicator): A Boolean value (true/false) indicating whether significant atomic clashes are present in the predicted structure. If more than 50% of the atoms in a given chain experience collisions or if a chain contains over 100 colliding atoms, the value is set to true, indicating potential structural issues.

a. ipTM (Interface pTM-score): Ranges from 0 to 1, representing the interface TM-score, which assesses the reliability of intermolecular interactions. Larger values indicate greater confidence in predicted interactions between different chains.
b. Ranking_Score: Ranges from −100 to 1.5, used to rank predicted structural models. A larger score corresponds to increased reliability of the structure.

**Table 2.** Summary of confidence scores for each structure originally downloaded from the AlphaFold3 server with four zinc ions added per monomer.

| Scores<br>ZIP | Fraction_Disordered | Has_Clash | ipTM | pTM | Ranking_Score |
| --- | --- | --- | --- | --- | --- |
| 1 | 0.18 | 0.0 | 0.59 | 0.62 | 0.68 |
| 2 | 0.16 | 0.0 | 0.68 | 0.72 | 0.77 |
| 3 | 0.18 | 0.0 | 0.59 | 0.63 | 0.69 |
| 4 | 0.16 | 0.0 | 0.52 | 0.53 | 0.6 |
| 5 | 0.33 | 0.0 | 0.52 | 0.53 | 0.69 |
| 6 | 0.49 | 0.0 | 0.44 | 0.45 | 0.69 |
| 7 | 0.43 | 0.0 | 0.56 | 0.57 | 0.78 |
| 8 | 0.23 | 0.0 | 0.56 | 0.58 | 0.68 |
| 9 | 0.12 | 0.0 | 0.73 | 0.74 | 0.79 |
| 10 | 0.5 | 0.0 | 0.43 | 0.43 | 0.68 |
| 11 | 0.22 | 0.0 | 0.7 | 0.71 | 0.82 |
| 12 | 0.22 | 0.0 | 0.45 | 0.46 | 0.56 |
| 13 | 0.25 | 0.0 | 0.59 | 0.63 | 0.72 |
| 14 | 0.24 | 0.0 | 0.54 | 0.56 | 0.66 |

These metrics provide a quantitative assessment of the confidence into the structures across different regions, allowing for a more comprehensive evaluation of the predicted models.

For certain members with lower confidence scores, structural optimization is particularly critical. In the ZIP family, the loop regions, N-terminus, and C-terminus are positioned outside the membrane region in the extracellular, and also intracellular regions [12]. These fundamental constraints must be adhered to during subsequent structural optimization to ensure biologically accurate modeling.

### 2.2 Optimization

The initial models obtained from AlphaFold3 often exhibit flexible loops connecting the TM α-helices, which are unlikely to reflect the biological conformation. Therefore, we assessed the feasibility of these models and performed further structural optimization to correct any unrealistic features. To obtain highly reliable protein structures, the strategy of homology modeling was employed to address regions of low credibility within the initial structural prediction using Modeller version 9.17 [34]. This approach was particularly focused on resolving uncertainties at the C-terminal and N-terminal regions, where confidence levels in the predicted structures were notably lower. By leveraging homology modeling, it was possible to incorporate information from structurally similar proteins, thereby enhancing the overall accuracy and reliability of the modeled structures. Importantly, all the structural templates used in this process were derived from the initial protein models predicted by AlphaFold3, which served as the foundational framework for further refinement. This combination of state-of-the-art prediction tools and traditional modeling techniques allowed for the construction of highly plausible and biologically relevant protein structures. VMD was used to visualize the structures [35]. After model construction, energy calculations (e.g., molecular force fields) and scoring functions like DOPE [36] or GA341 [37] were used to evaluate structural plausibility. For scoring functions, the DOPE potential was calculated with a grid spacing of 0.5 Å and a dielectric constant of ε=4, while GA341 scores incorporated secondary structure compatibility and residue-residue contact metrics. Validation thresholds were set as follows: Ramachandran Plot analysis required >90% of residues in allowed regions [38], ProSA Z-scores within ±2 standard deviations of native structures [39], and Verify3D 3D-1D profile scores >0.2 [40]. QMEANDisCo45 was additionally applied to evaluate global model quality, with scores >0.6 considered acceptable [41]. In the process of homology modeling, we fixed the high credibility region structure, such as the ECD and the TMD. After the adjusted protein structures were obtained, the conjugate gradient method within the GROMACS molecular dynamics simulation package was used to optimize the structures [42]. The simulations were performed under explicit solvent conditions using the TIP3P water model, with a cubic periodic boundary box extending 12 Å beyond the protein surface. The system was neutralized with counterions (Na^+^/Cl^−^) to achieve a physiological ion concentration of 0.1 M. The CHARMM36m forcefield was used to describe the interactions of the protein monomers [43]. The cut-off for the nonbonded interactions was set to 12 Å, and the particle mesh Ewald (PME) algorithm was applied to treat the long-range electrostatic interactions.

## 3. Results

The predicted structures are presented here according to their similarity based on sequence homology (Figures 1 and 2). In the ZIP family, dimerization is driven by the leucine zipper motif, characterized by a heptad repeat pattern [41–43]. Hydrophobic residues at every seventh position form an amphipathic α-helix, enabling two α-helices to interact via hydrophobic interactions and to form a coiled-coil dimer [44–46]. In addition, in ZIP family members with an ECD, interactions in this domain have a role in dimerization as well. Thus, the conserved ‘PAL’ motif in the ECD stabilizes the dimer. In hZIP4, the ‘PAL’ motif at the dimerization interface, consisting of residues SER275, PRO276, LEU278, and 279, plays a critical role in dimer formation [12].

**Figure 1.**
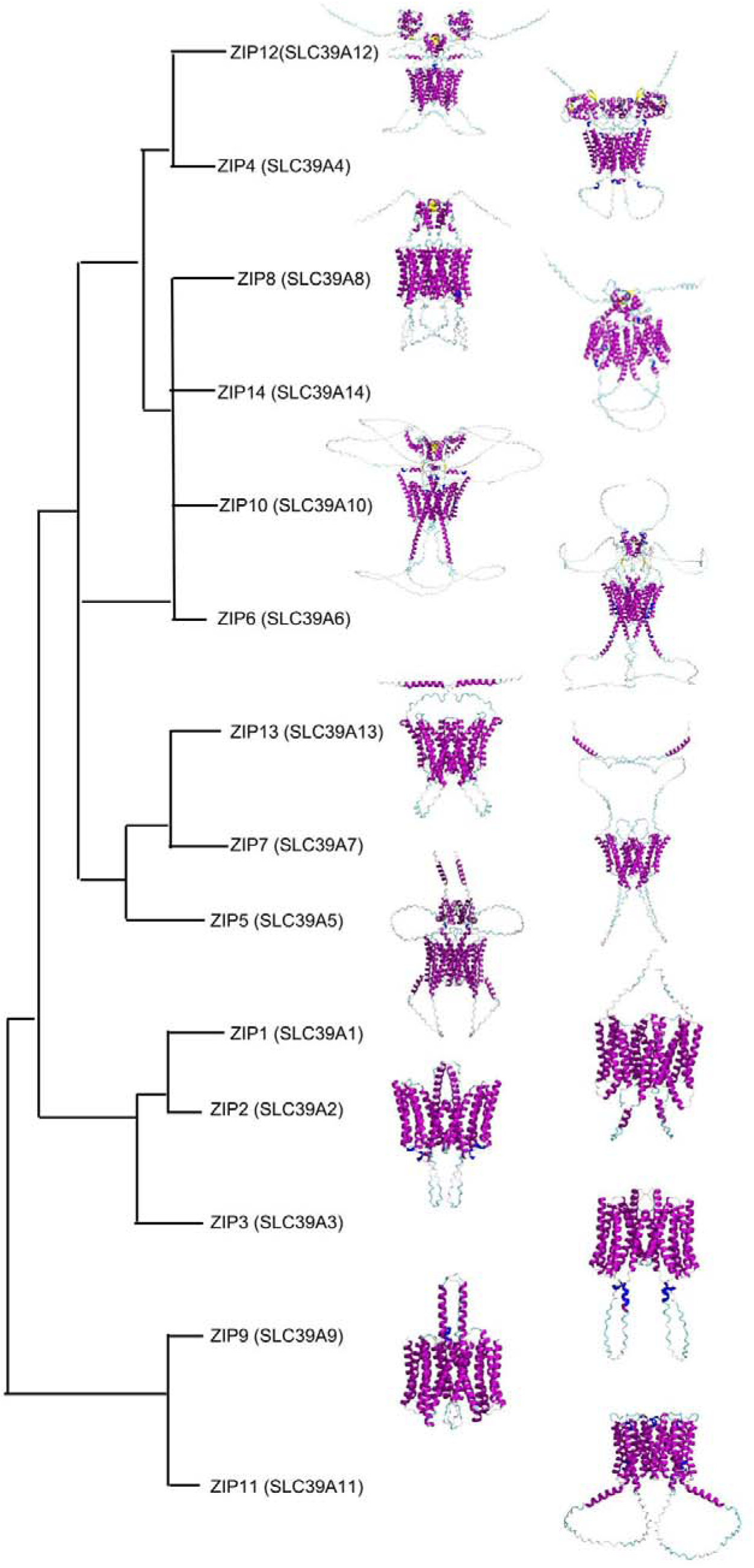
Summary of the 3D structures of all 14 human ZIP members - with secondary structure emphasized - following the sequence similarity. Purple are α-helices. Yellow are β-sheets. Light green are loops. The LIV1 subgroup contains ZIP4-8, 10, and 12-14. Among them, the structures of ZIP4,12,8, and 14 are similar while the structures of ZIP5,7, and 13 are similar to each other, too. ZIP4,12,8,14, and 5 all have an ECD. ZIP7 and 13 have long N-termini. The other three groups contain ZIP1,2,3,9, and 11, which do not have an ECD. Accordingly, their structures are smaller than those of the members of the LIV1 subfamily.

**Figure 2.**
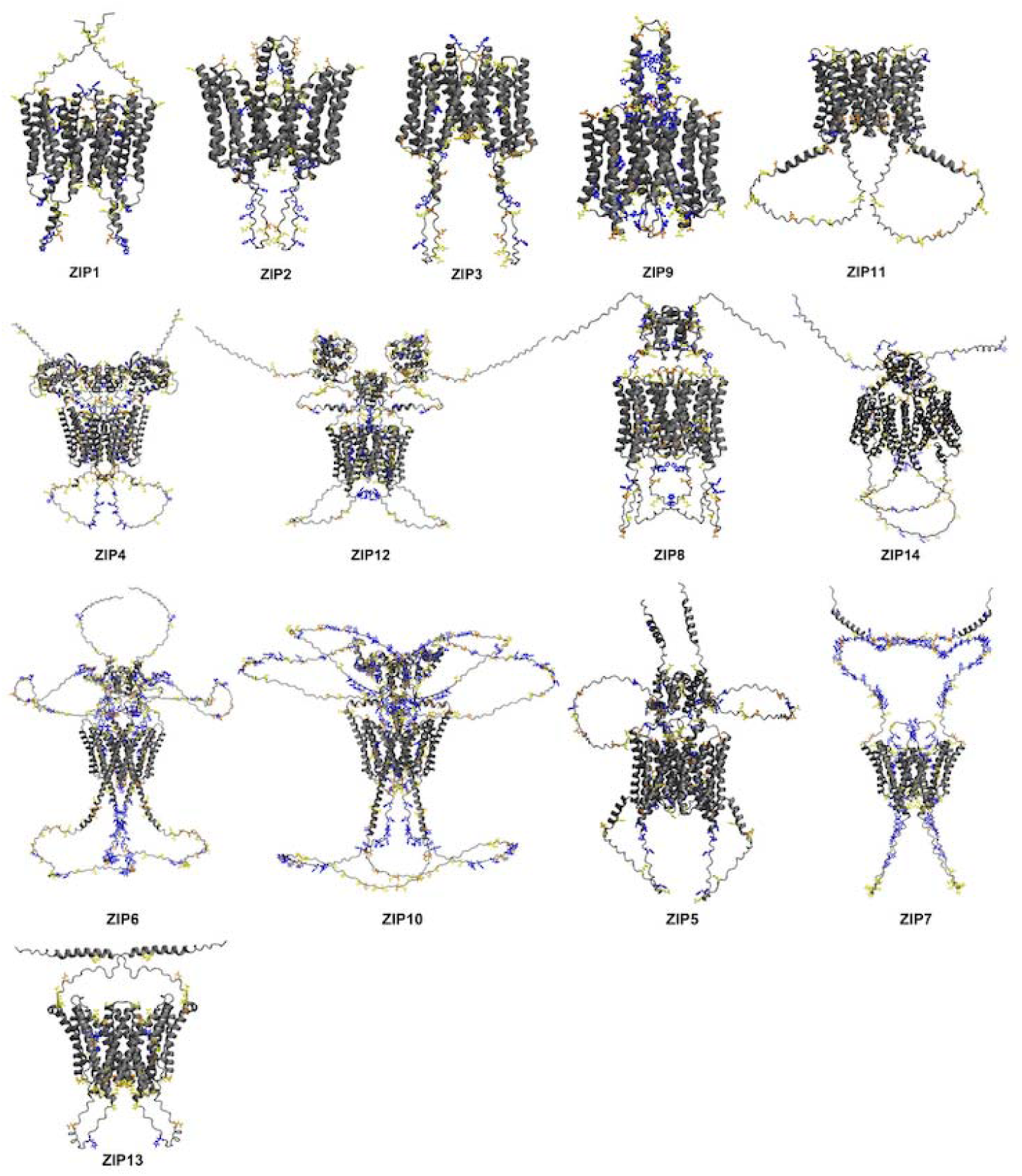
The overall dimeric structures of all 14 human ZIP members and the distribution of HIS, ASP, and GLU residues. Blue represents HIS, yellow GLU, and orange ASP residues. ZIP1,2,3,9, and 11 do not have ECDs. ZIP4 and 12 have 2 ECDs in each monomer. ZIP8,14,6,10, and 5 only have one ECD in the monomer. The ECD contains many HIS and GLU residues that may attract zinc ions. Except ZIP9, all the other 13 members have a very long loop in the cytoplasmic domain between TM3 and 4. ZIP7 has a very long N-terminal that contains HIS motifs. ZIP6 and 10 have the most complicated structures. In their ECDs, there are long loops between the α-helices containing lots of HIS and ASP motifs.

### 3.1 The transmembrane domain (TMD)

Each ZIP member forms a dimer, and each TMD monomer is composed of eight α-helices, which are connected by loops that can be categorized based on their location (Figure 2). Extracellular loops are found between TM2-TM3, TM4-TM5, and TM6-TM7, while cytoplasmic loops connect TM1-TM2, TM3-TM4, TM5-TM6, and TM7-TM8. Both N- and C-termini face outwards. For all members, the eight α-helices of one monomer can be divided into two four-helix bundles. HB1 contains TM1,4,5, and 6 and HB2 has TM2,3,7, and 8 (Figure 3, A and C). The zinc ion transport channel is between these bundles and is composed of TM2,7,4, and 5 (Figure 3, B and D) [19]. For HB1, TM4 and 5 are positioned on one side, while TM1 and 6 are on the opposite side. These four α-helices do not intersect (Figure 3, A and C). HB2 participates in the dimerization. In ZIP1,2,3,5,7,9,11, and 13, HB2 adopts a cross-like arrangement, with TM8 intersecting with TM2, and TM7 intersecting with TM3. TM8 and 7 are almost parallel (Figure 3, A). In ZIP4,8,6,10,12, and 14, TM3 is positioned on top of TM7, aligning with TM8 in a nearly parallel orientation. This configuration places TM3 and 8 at the forefront, followed by TM2 and 7 in a cross pattern (Figure 3, C).

**Figure 3.**
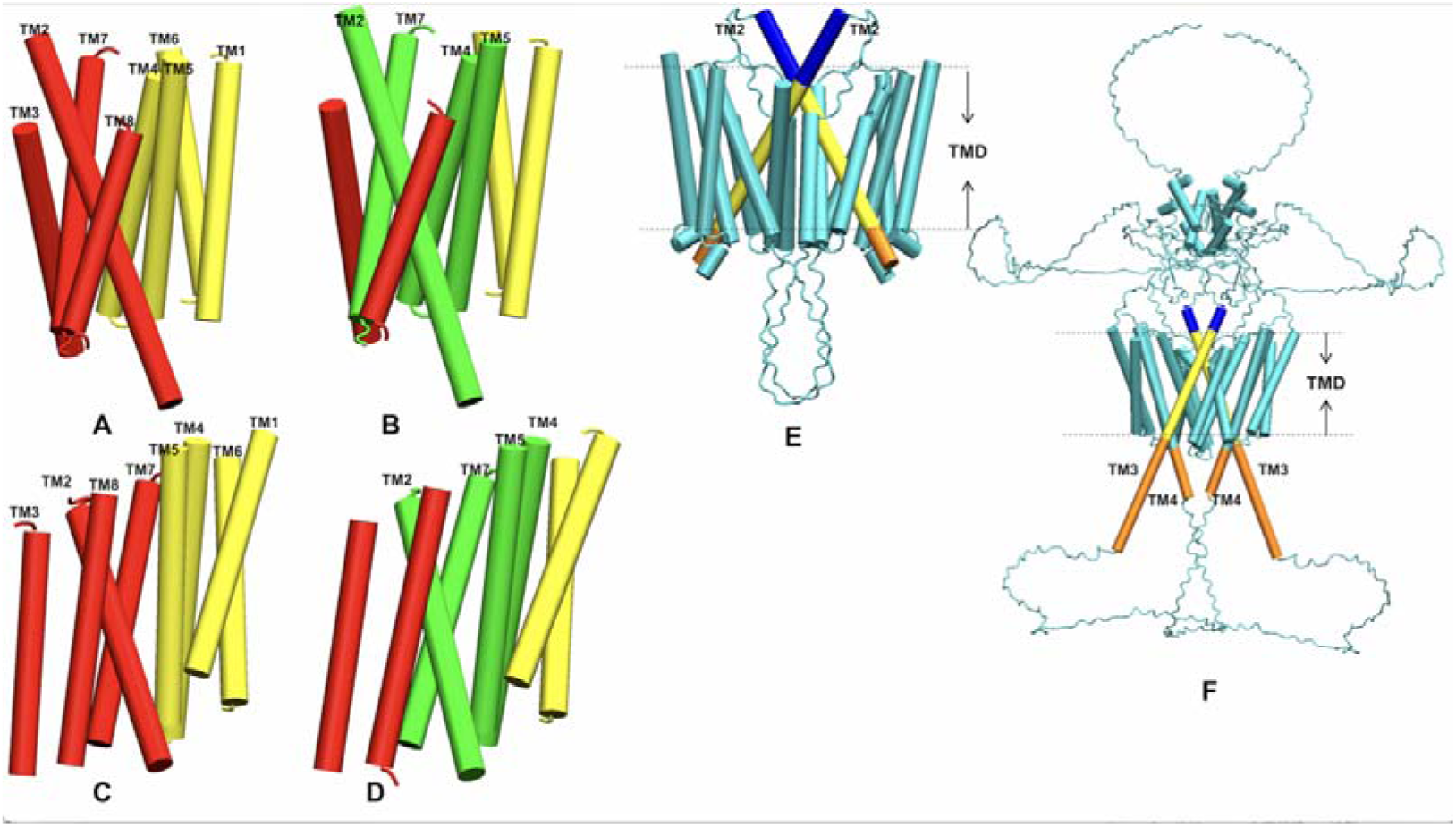
The arrangement of α-helices in the TMD monomer. Yellow denotes HB1 containing TM1,4,5, and 6. Red denotes HB2 containing TM2,3,7, and 8. The metal ion transport cavity is shown in green and is composed of TM2,7,4, and 5. TM4 and 5 belong to HB1 while TM2 and 7 belong to HB2. A and B represent the TMDs from ZIP1,2,3,9,11,5,7, and 13. A shows the locations of the two bundles. B shows the transport channel. C and D show the arrangement of α-helices in the TMDs of ZIP4,12,8,14,6, and 10. C shows the locations of HB1 and HB2. D also shows the transport channel. Compared with B, the locations of TM3 and 8 in D are different. E and F show the longest α-helix of the TMD. Blue is the part in the ECD. The part within the membrane is shown in yellow, the cytoplasmic domain in orange. E shows the location of the longest TM2. F shows the location of the longest TM3. Although TM4 is not the longest, it still is partially in the cytoplasmic domain.

There are α-helices extending beyond the TMD. Notably, ZIP1,2,3,9, and 11 have a relatively long TM2, containing 45, 51, 44, 52, and 34 residues, respectively (Figure 3, E). In ZIP9, TM4 also contains about 5 residues in the cytoplasmic domain. ZIP6,10,12, and 14 have longer TM3 segments (Figure 3, F), with lengths of 68, 69, 38, and 39 residues, respectively. In ZIP7 and 13, TM4 is the longest, measuring 42 and 39 residues, respectively. has TM8 and 4 of equal length, each containing 34 residues. In contrast, ZIP4 and 8 do not have elongated α-helices.

### 3.2 The extracellular domain (ECD)

The ECD of ZIP4 in the monomer consists of two distinct subdomains: the N-terminal histidine-rich domain (HRD) and the C-terminal PAL-containing domain (PCD) [12]. Only members of the LIV1 subfamily possess an ECD. For instance, ZIP4 features both the HRD, and the PCD while subgroups ZIPI and ZIP II contain only the PCD. The dimerization of the ECD is facilitated by domain swapping of the PCDs between the two monomers. Within this interface, the conserved PAL motif is crucial, as its residues are positioned at the intersection of two α-helices at the core of the PCD dimer [12,47]. For all members of the ZIP family, the ECD includes the N-terminus, C-terminus, and loops connecting α-helices in the TMD, and in some members long α-helices in the TMD extend beyond the TMD (Figure 4). Additionally, in ZIP4,5,6,8,10,12, and 14, the ECD also includes more complex arrangements with α-helices, loops, and β-sheets (Figure 5) (S, Figure 1).

**Figure 4.**
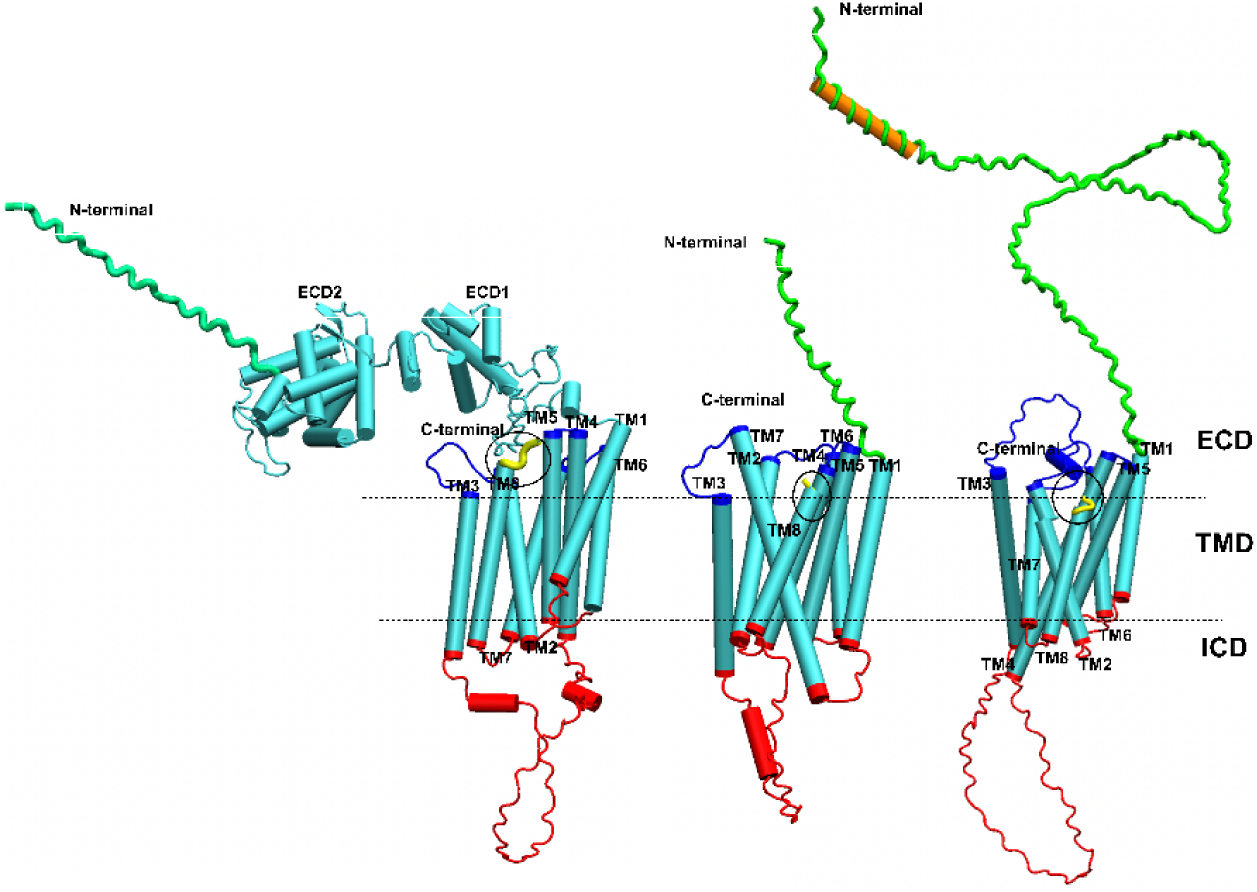
The locations of the ECD, TMD and cytoplasmic domain (ICD). Shown from left to right, are ZIP4, 1, and7, respectively. The figure also shows the locations of the N-terminus (green), C-terminus (yellow with black circles) and loops (blue or red).

**Figure 5.**
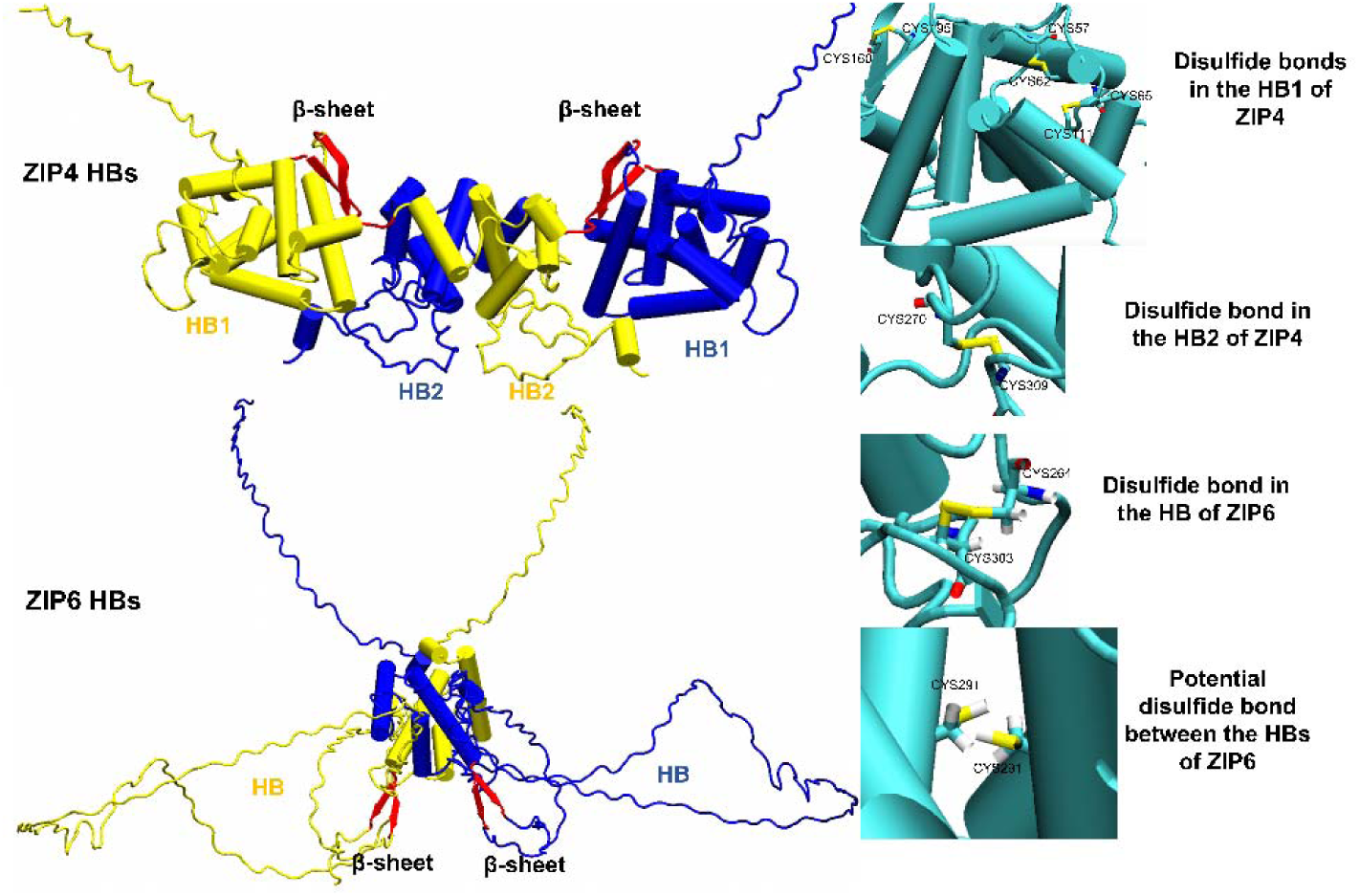
The ECDs in ZIP4,12,8,14,6,10, and 5. ZIP4 and 12 have two ECDs in the monomer while ZIP8,14,6,10, and 5 have only one ECD in the monomer. The ECD of the first monomer is in blue, and the ECD of the other monomer in yellow; β-sheets are in red. On the right, the disulfide bonds between the α-helices are shown.

Only ZIP5,7, and 13 have very long N-termini but the N-termini adopt an α-helical structure (Figure 6). The length of the N-terminus varies across ZIPs, ranging from 25 to 132 residues, with ZIP7 having the longest. In ZIP5,7,10, and 13, the extended N-terminus also forms a single α-helical structure at the beginning of the N-terminus. The C-termini are generally short, from a single (ZIP1,5, and 11) to ten residues (ZIP6) (Figure 6).

**Figure 6.**
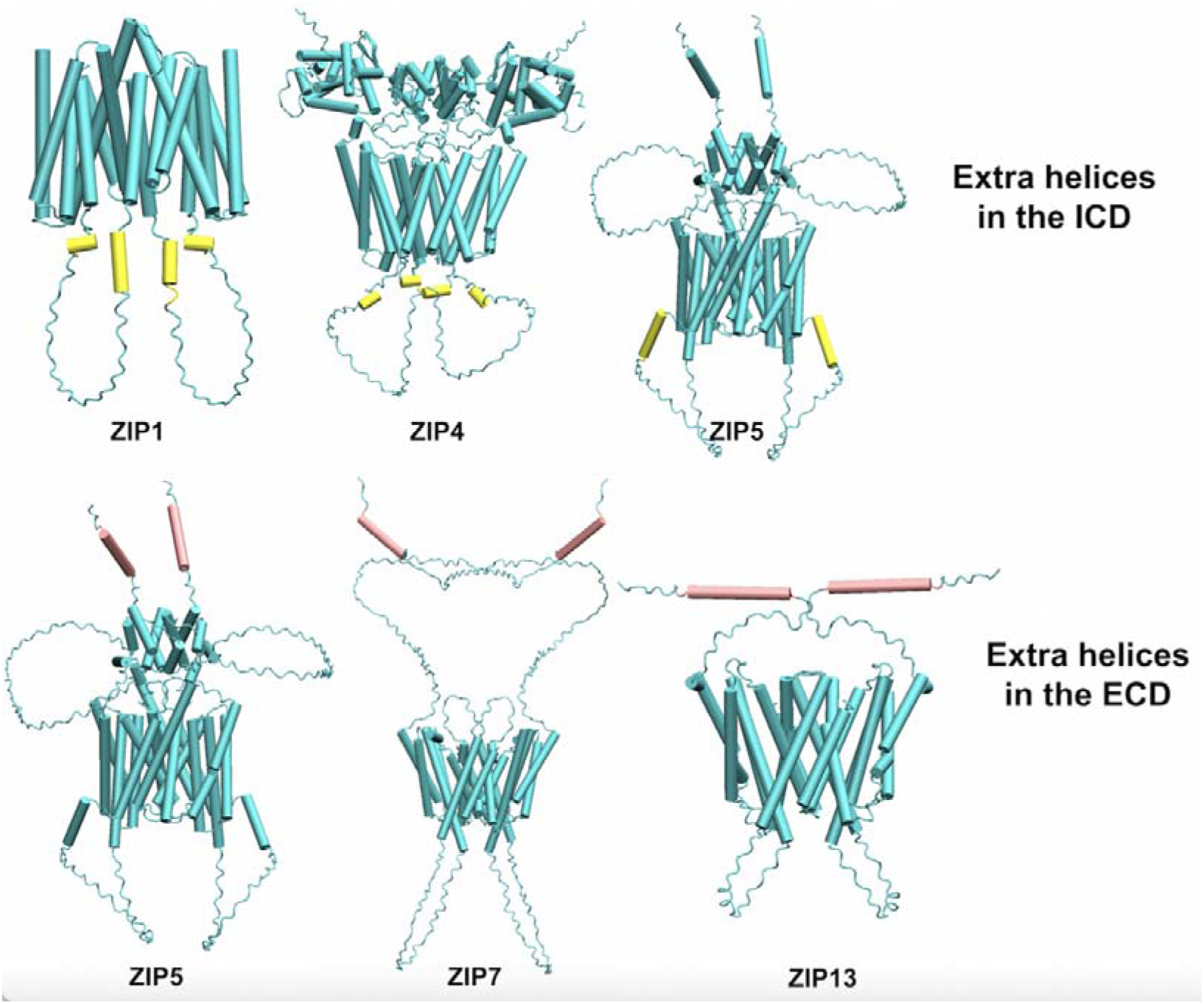
The extra α-helices between the loops in the ECD (pink) and the ICD (yellow). ZIP5,7, and 13 have α-helices in the ECD, while ZIP1-4,11, and 14 have α-helices in the ICD. ZIP5 has both extracellular and cytoplasmic α-helices.

The ECDs of ZIP4,5,6,8,10,12, and 14 have more complex structures (Figures 1 and 5). The α-helices are connected by loops, β-sheets, and disulfide bonds. The monomers of ZIP4 and 12 feature ECDs composed of two distinct regions: ECD1 and ECD2. ECD2 from one monomer forms a dimer with the ECD2 of another monomer. The number of α-helices in the ECD varies across different ZIP members. For example, in ZIP4, ECD1 contains 9 α-helices and ECD2 contains 4 α-helices. In ZIP12, the ECD1 contains 8 α-helices, while ECD2 contains 5. The number of α-helices in the ECDs of the monomer is 6 in ZIP5, 5 in ZIP8, 3 in ZIP6, 8 in ZIP10, and 5 in ZIP14. Notably, ZIP6 and 10 possess exceptionally long loop structures within their ECDs. In ZIP6, the loop between the second and third α-helix comprises 204 residues, while in ZIP10, a loop of 199 residues connects the fourth and fifth α-helix.

The β-sheets also help maintaining the structure of the ECD (Figure 5). ZIP members have ECDs that contain β-sheets forming a hydrophobic core [48]. These sheets are found in ZIP4,6,8,10,12, and 14, connecting α-helices within the ECD and maintaining the structure. For example, in ZIP4, ECD1 and 2 are connected by a β-sheet. ZIP6 has a β-sheet connecting the ECD and the TMD. In particular, the mammalian ZIP4 ECD has been shown to contain an eight-stranded β-sheet fold [48,49].

### 3.3 The intracellular domain (ICD)

The ICD consists of two distinct structural components. The first component comprises loops connecting TMDs, including the loops between TM1–TM2, TM3–TM4, TM5–TM6, and TM7–TM8 (Figure 4). Except for ZIP9, the TM3–TM4 loop is very long and is called IL2 [45]. ZIP6 and 10 have the longest TM3–TM4 loops, containing 100 and 117 residues, respectively. This loop is functionally significant, as it contains lots of HIS and GLU residues (Tables 3,4, and 5) [50].

In the ICD, there are also extra α-helices between the loops of α-helices of the TMD. For example, ZIP2 has two α-helices of four residues each on the TM1-TM2 loop. On the TM3-TM4 loop, ZIP3 has two additional α-helices, and similarly, ZIP4 and 14 each form two small α-helices. ZIP7 forms an α-helix with 8 residues between TM6 and 7 (Figure 5). There are elongated TM2 helices in ZIP1,2, and 3, as well as TM3 in ZIP6 and 10 (Figure 6).

### 3.4 Distribution of sequence motifs with potential zinc-binding residues

#### 3.4.1 Cysteine residues (CYS)

Disulfide bonds are critical for maintaining the stability of some ZIP proteins [51]. In the ECD, ZIP13’s N-terminus contains cysteine residues (CYS4, 6, and 9), where disulfide bonds may form between CYS4 and 6 though the distance between these two sulfur atoms is 4.97 Å and thus beyond the S-S bond length of about 2 Å. In ZIP proteins with ECDs, disulfide bonds are crucial for stabilizing monomers and dimers. ZIP4 and 12 have intramolecular disulfide bonds to help maintaining monomer stability, while ZIP8,14,6,10, and 5 have both intramolecular and intermolecular disulfide bonds for both dimer and monomer reinforcement. For instance, ZIP4 has three key disulfide bonds in ECD1. The first is CYS57-CYS62. The distance between the two S atoms is 1.84 Å. The second is CYS65-CYS111 with an S-S distance of 1.85 Å, and the third is CYS160-CYS195, which is also on the β-sheet, with a distance of 1.75 Å. There is also one in ECD2 (CYS270-CYS309) and the distance is 2.01 Å. ZIP8 has disulfide bonds in both monomer and dimer. The CYS101-CYS101 (distance of 2.21 Å) between the fourth helices of each monomer to connect the dimer, while the CYS74-CYS113 bond helps maintaining the structure of the monomer (distance of 2.07 Å) (Figure 5) (S, Figure 1).

In ZIP1,2,3, and 9, cysteine residues are distributed across the TMD, where some are positioned to potentially form disulfide bonds. For example, CYS181 on TM4 and CYS242 on TM6 are close, but not sufficiently close to form a disulfide bond. In ZIP1, the distance between these 2S atoms is 7.84 Å). There are more CYS residues in ZIP2 than in ZIP1,3, and 9. At the bottom of TM3, there is a CC motif with CYS126 and 127. In ZIP13, CYS residues are distributed at the top of the TMD α-helix. In the ICD, ZIP8 has CSC (CYS374, SER375, and CYS376 on TM6) and CTC (CYS289, THR290, and CYS291) motifs on the loop between TM3 and 4 (Figure 7). Although these CYS motifs do not form a zinc ion binding site, CYS residues are also important for the zinc ion transport mechanism. For the zinc ion binding sites in the ZIP8, CYS310 is one of the ligands forming a potential M site (M3 site). In ZIP1, the exit site (M_ex_) contains CYS74.

**Figure 7.**
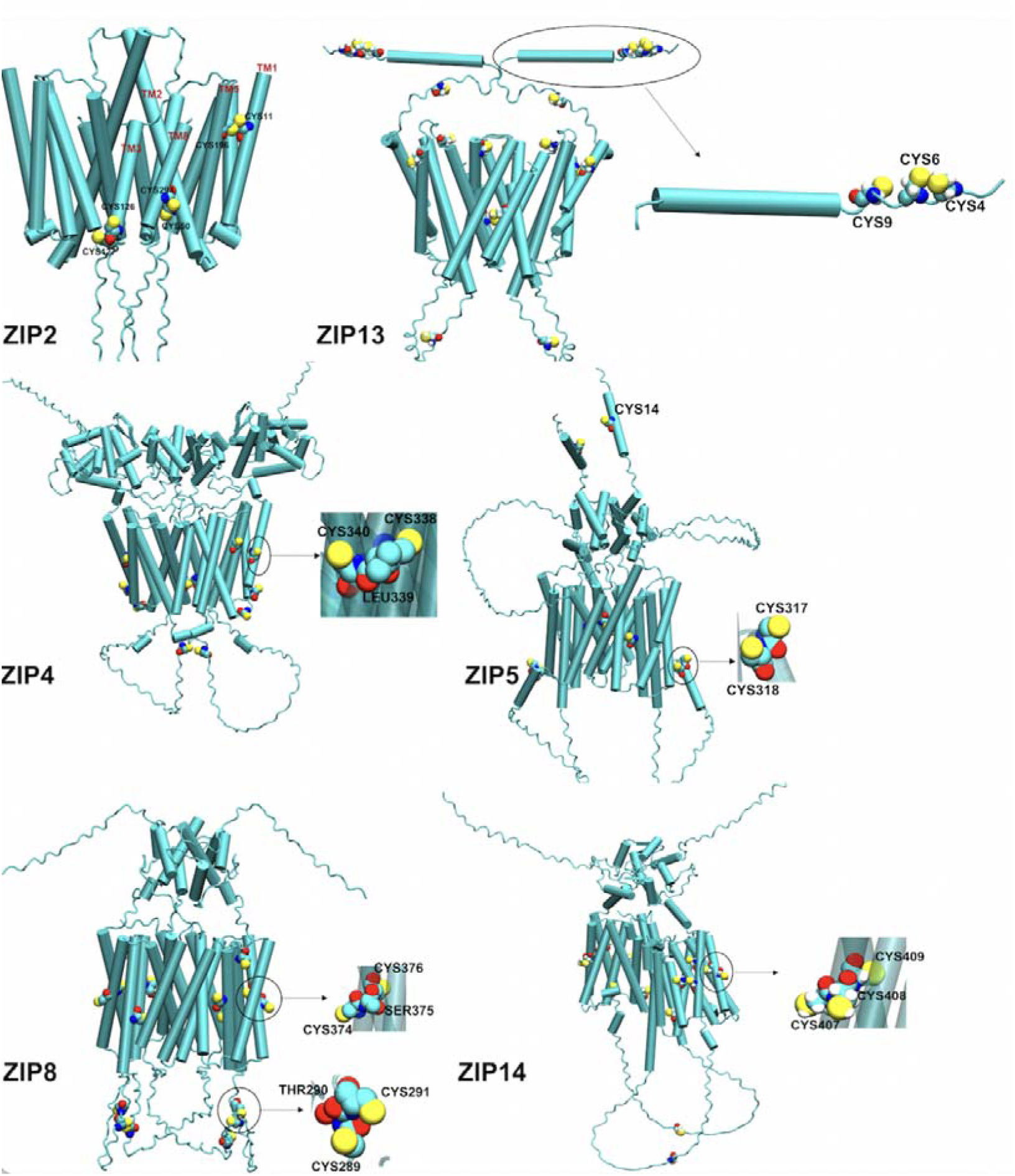
Distribution of CYS residues. ZIP2 has a CC motif on TM3. ZIP13 has 3 CYS residues on the N-terminal. ZIP4 has a CLC motif on TM1. ZIP5 has a CC motif on the extra α-helix in the ICD. ZIP8 has CSC and CTC motifs. ZIP14 has a CCC motif on TM6.

#### 3.4.2 Glutamic acid residues (GLU)

The GLU-rich motifs in human ZIP transporters are summarized in Table 3.

**Table 3.** The distribution of glutamic acid residue motifs.

|  | **Extracellular domain**<br>**(ECD)** | **Transmembrane domain**<br>**(TMD)** | **Intracellular domain**<br>**(ICD)** |
| --- | --- | --- | --- |
| ZIP9 | EHE |  |  |
| ZIP11 |  |  | ESE |
| ZIP4 | EPE | SEE | EES |
| ZIP12 | REE, QEE, EKESE | CEE |  |
| ZIP8 | EIELE | AEE, CEE |  |
| ZIP14 |  | ENEENE, EEG, CEE | EDE, QEE |
| ZIP6 | EHE, EES | REE | NEE, EEEE |
| ZIP10 | HEEH, ELE, EKE, EGE | EEY | ELE, EEEE, ETE, EEH, EES |
| ZIP5 |  | EE, EQE |  |
| ZIP7 |  | EEE | EEEEKE, EES |
| ZIP13 | ESE | EEED |  |

In the LIV1 subfamily of ZIP proteins, a unique “metalloproteinase” type of motif (HEXPHEXGD) contains conserved GLU residues critical for zinc binding and transport across the membrane [18]. However, ZIP8 and 14 have an EEXPHEXGD motif instead. The GLU residues form the M binding sites in the TMD. The specific arrangement of GLU residues could dictate substrate specificity in the ECD, too, e.g., the ZIP members with the ECD have more GLU motifs. For instance, ZIP12 has REE, QEE, and EKESE motifs, and ZIP10 has HEEH, ELE, EKE, and EGE motifs. In the TMD, all the members of LIV1 have GLU motifs, especially EE motifs like SEE or EEY. In the ICD, ZIP10 has lots of GLU motifs such as ELE, EEEE, ETE, EEH, and EES. In ZIP9, the EHE motif belongs to the HEHEHSHDH motif, which is on the loop between TM2 and 3 in the ECD. In ZIP1, 2, 3, 9, and 11, GLU residues form zinc ion binding sites. For example, the entrance sites (M_en_ sites) of ZIP2 and 3, which are on the top of the TMD, contain only GLU residues while ZIP4 does not have GLU residues around any zinc ion binding sites. ZIP12 has GLU648 at the M_ex_ site, which is at the bottom of the TMD. The two GLU residues from the CEE (CYS342, GLU343, GLU344) motif are part of the M1 site in Zip8 together with HIS347. Also, in ZIP14, GLU377 from the CEE motif is part of the M1 site together with ASN348 and ASP351. For ZIP13, GLU128 is in the M_en_ site and binds the zinc ion for transport to the M1 site. In the other binding sites of ZIP13, there are also GLU residues although they are not forming a specific motif. GLU530 from the EE motif on TM8 of ZIP5 forms the M_en_ site with HIS264 and 268. For the M_ex_ site of ZIP5, GLU428 together with ASP391, ASN395 and HIS427 may be involved in zinc ion binding (Table 4).

**Table 4.** The metal binding sites in the different ZIP members. All the ligand donor atoms are within 3 Å of the zinc ion in the TMD.

| **Name** | **M<sub>en</sub> Site** | **M1 Site** | **M2 Site** | **Potential M Sites** | **M<sub>ex</sub> Site** |
| --- | --- | --- | --- | --- | --- |
| ZIP12 | HIS419, HIS428, GLU686 | HIS551, ASP555, HIS580 | ASP548, HIS584, ASP588 |  | GLU648 |
| ZIP4 | HIS388, HIS390, ASP643 | ASP375, HIS507, HIS536 | ASP504, HIS540, HIS536, ASP544 |  | HIS358, ASP614 |
| ZIP8 | ASP189, ASP193 | GLU343, GLU344, HIS347 | ASP311, ASN315, GLU344 | 1. ASP311, HIS34, ASP351<br>2. CYS310, HIS314, ASP410 |  |
| ZIP14 |  | ASN348, ASP351, GLU377 | HIS347, GLU376, HIS380 | 1. ASP344, HIS38, ASP384<br>2. HIS347, HIS380, ASP443 |  |
| ZIP10 | ASP454, HIS458, HIS709 | HIS680, GLU710, HIS713 | ASP677, ASN681, GLU710 | ASP677, GLU714 | ASP787, ASP789, HIS793 |
| ZIP6 | 1. HIS380, HIS385, GLU746<br>2. ASP372, HIS376, HIS635 | HIS606, ASN607, ASP610, GLU636 | ASP603, GLU64, ASP643 |  |  |
| ZIP13 | GLU128, ASP371 | ASP228, ASN229, HIS232, GLU258 | ASN225, GLU262, ASP265, GLN283 | ASN120, ASP228, GLU258, HIS261 |  |
| ZIP7 | ASP183, HIS187, HIS329 | HIS329, ASN330, ASP333, GLU359 | ASP326, HIS362, GLU363, ASP366 |  |  |
| ZIP5 | HIS264, HIS268, GLU530 | HIS394, ASP398, HIS423 | ASP391, HIS427 |  |  |
| ZIP1 | ASP91, HIS190, GLU276 | ASP87, HIS190, GLU194, HIS217 |  |  | SER70, CYS74, GLU295 |
| ZIP2 | GLU67, GLU71,<br>GLU262 | HIS63, HIS175,<br>GLU179, HIS202 |  |  | HIS216,<br>GLU276,<br>GLU281 |
| ZIP3 | GLU69, GLU193 | HIS180,<br>GLU184, HIS207,<br>GLU208 |  |  | GLU285,<br>GLU288 |
| ZIP9 | GLU57, HIS60,<br>HIS305, HIS307 | HIS155, ASP159,<br>HIS185 |  |  | ASP123,<br>HIS261 |
| ZIP11 |  | HIS204, GLU208,<br>GLN240, GLU244 | ASN205, GLU208,<br>ASN241, GLU244 |  | ASP96,<br>ASP308,<br>ASP309 |

#### 3.4.3 Aspartic acid residues (ASP)

ASP residues are distributed across different domains of the ZIP proteins (Table 5) and have roles in metal ion transport [52]. The DD motif is the most prevalent ASP motif. In ZIP1,2,3,9, and 11, ASP residues are sparse. ASP309 of ZIP11 forms the M_ex_ site from a DD motif with ASP96. In ZIP members with an ECD, ASP residues are primarily found on surface regions, except for ZIP6 and 10, where they cluster on long loops. ZIP4,12,8, and 14 have ASP-enriched ECDs, though distinct ASP motifs are rare. In contrast, ZIP6 and 10 feature specific motifs like DHD and DLDPD in long loops of the ECD. Within the TMD, ASP residues are mainly located in the zinc transport cavity between TM4 and 5, where they often are part of the metal-binding sites (Table 4). In ZIP4-8,10,12,13 and 14, typically, three ASP residues are between TM4 and 5, with one located on TM5 (Figure 8). In ZIP12, ASP555 on TM4 together with two HIS residues form the M1 site. ASP548 on TM4 and ASP588 on TM5 with HIS584 form the M2 site. In the ICD, ASP residues are mainly located on the loops between TM3 and 4. ASP643 from a DD motif in ZIP4 with HIS388, THR389 and HIS390 (HTH) may form an M_en_ binding site (Table 4).

**Figure 8.**
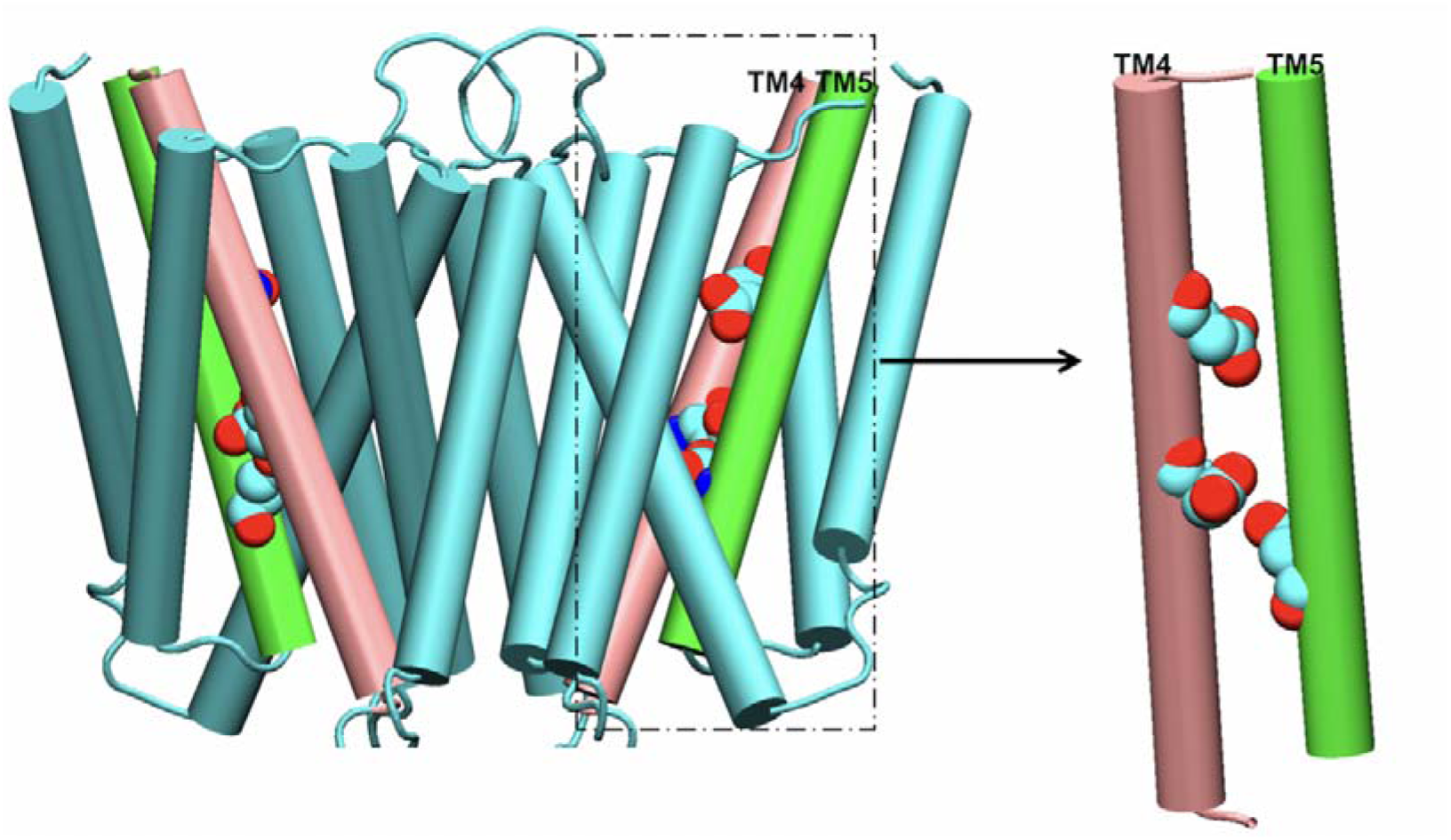
In ZIP4-8,10,12,13, and 14, three aspartic acid (ASP) residues are between TM4 (pink) and 5 (green). The ASP residue on TM5 is part of the transport site (Table 5).

**Table 5.** The distribution of aspartic acid residue motifs.

|  | **Extracellular domain**<br>**(ECD)** | **Transmembrane domain**<br>**(TMD)** | **Intracellular domain**<br>**(ICD)** |
| --- | --- | --- | --- |
| ZIP2 | DAD |  |  |
| ZIP9 | 2DD |  | DD |
| ZIP11 |  | DD |  |
| ZIP14 |  |  | DLD |
| ZIP6 | 2DHD, DHDSD |  | 2DD, DDD |
| ZIP10 | 2DD, DLDPD |  | DD, DGD, DSD |
| ZIP7 | 2DD, DHD, DLD |  |  |

#### 3.4.4 Histidine residues (HIS)

The HIS-containing motifs are summarized in Table 6.

**Table 6.**
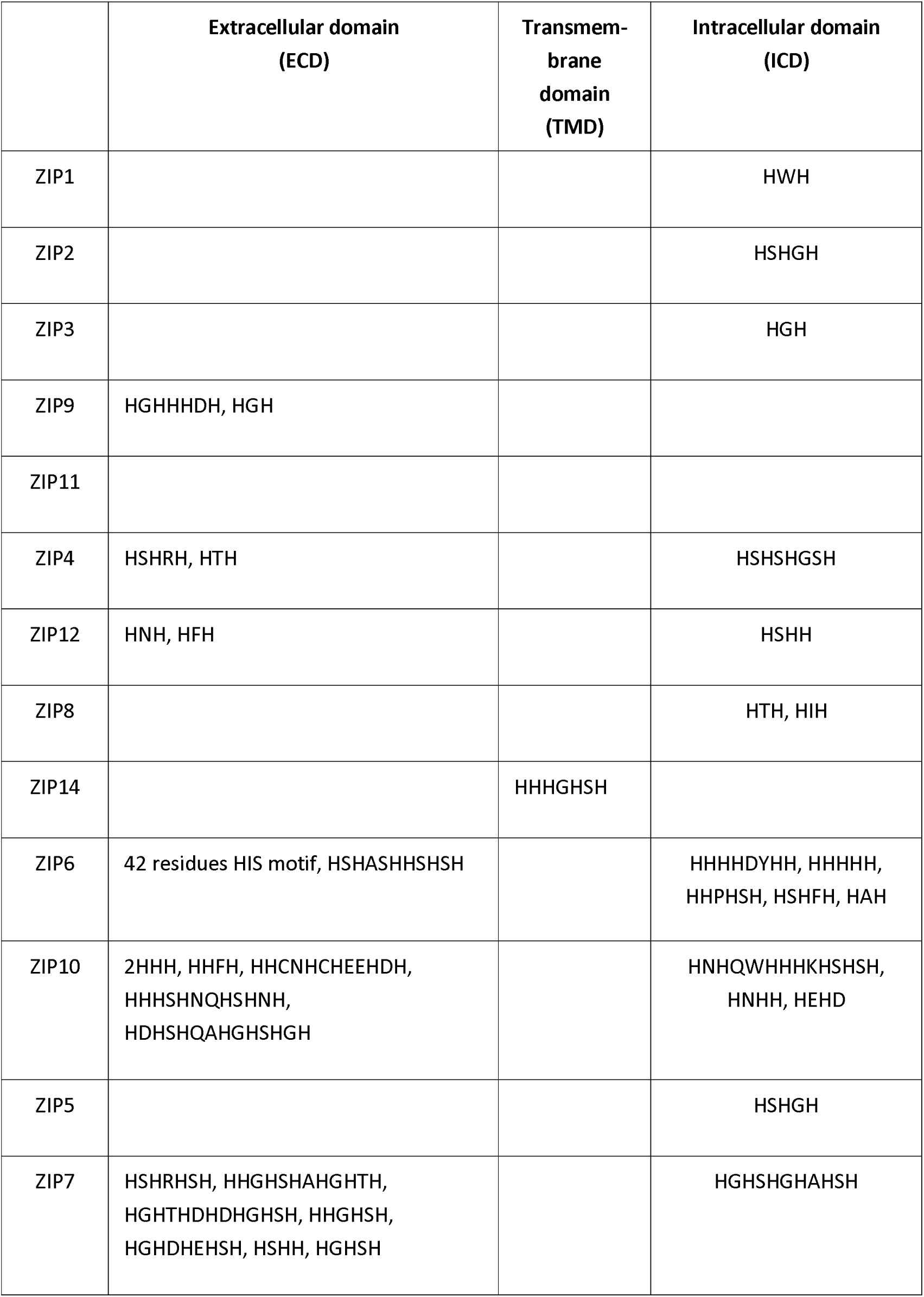
The distribution of histidine residue motifs.

ZIP4 has a HIS-rich loop in its ECD that binds the zinc ion with micromolar affinity [12]. ZIP6 and 10 contain HIS motifs that are essential for zinc ion binding and transport [18,53]. In the ECD, many ZIP proteins have HIS-rich regions. In ZIP1, the IL2 loop contains an HWH motif composed of HIS158 and 160, which is essential for zinc transport [4]. Similarly, ZIP2 features an HSHGH motif in its IL2 loop, highlighting the significance of HIS residues in this region [2]. ZIP9,4,12,6,10, and 7 have HIS motifs in the ECD. Among them, ZIP6,10, and 7 have more His-containing motifs. The loop between the second and third α-helix of the ECD monomer includes a motif composed of 42 residues (HIS93 to HIS134), containing 22 HIS residues along with ASP, SER, and GLU residues. In ZIP7, HIS residues are prevalent, particularly on the N-terminus, where motifs such as HSHRHSH (HIS45–HIS51), HHGHSHAHGHTH (HIS55–HIS68), HGHTHDHDHGHSH (HIS73–HIS85), HHGHSH (HIS89–HIS94), and HGHDHEHSH (HIS106–HIS114) are located. In the TMD, HIS residues are mainly in the transport channel. In ZIP1, HIS residues located between TM3 and 4 are required for zinc transport across the plasma membrane [54]. In ZIP14, an HHHGHSH motif is located at the base of TM3 within the TMD. The HIS residues in the ICD are mainly on the loop between TM3 and 4 (IL2). For the members of the LIV1 subfamily, there are HIS-containing motifs on this loop but close to the TMD. For example, ZIP4 has five HIS residues on the loop between TM3 and 4, alternating with four SER residues to form an HSHSHGSH motif (HIS438–HIS448) near the TMD. Similarly, ZIP12 has an HSHH motif (HIS486, SER487, HIS488 and 489) on the loop between TM3 and 4 near the TMD. By contrast, ZIP14’s TM3-TM4 loop contains only two HIS residues and does not form a specific motif. For ZIP6 and 10, the IL2 loop contains many HIS motifs. In ZIP6, in the loop between TM3 and 4, near the TMD, three motifs are identified: HHHHDYHH, HHHHH, and HHPHSH. In the same loop, further away from the TMD, additional motifs include HSHFH and HAH. ZIP10 has 3 motifs found on this loop, in which HNHQWHHHKHSHSH is near the TMD, and HNHH and HEHD are a bit further away from the TMD. In ZIP7, between TM3 and 4, motifs such as HGHSHGHAHSH and HGH are found. In ZIP7, HIS-rich motifs in the ICD play a role in zinc ion transfer from the TMD to the cytoplasm [53].

### 3.5 The proposed zinc ion transport pathways

Central to the function of ZIP transporters is a primary zinc-binding site, M1, the selectivity filter, but some ZIP proteins have a binuclear site containing another zinc-binding site, M2. In general, both sites are situated on TM4 and 5 of the TMD. However, in this study, TM2 and 7 are also observed to coordinate zinc ions at the M sites. For certain members, such as ZIP8 and 14, additional M3 and M4 sites have been identified. The M3 site comprises ligands located on TM4 and 5, whereas the M4 site includes ligands not only on TM4 and 5 but also on TM7.

All members of the LIV1 subfamily have a unique metalloproteinase-type motif that is involved in metal binding at the M1 site. Except for ZIP7 and 13, all other LIV1 subfamily members have a cysteine residue preceding the metalloproteinase motif, forming either a CHEXPHEXGD or CEEXPHEXGD motif. In ZIP4, HIS536 and 540 correspond to the first and second HIS residues of the HEXPHEXGD motif, forming the M1 and M2 sites, respectively, while in ZIP12, HIS580 and 584 form the M1 and M2 sites, respectively. In ZIP8, the first and second GLU residues (GLU343, GLU344) and the first HIS residue (HIS347) form the M1 site within the EEXPHEXGD motif. GLU344 also is part of the M2 site. GLU344 is the bridge residue between the M1 and M2 sites of ZIP8. In ZIP14, the second GLU residue of this motif (GLU377) forms the M1 site while GLU376 and HIS380 form the M2 site. HIS380 (the only HIS residue of EEXPHEXGD motif in ZIP14) also forms the M3 and M4 sites. In ZIP6, two GLU residues of this motif form the M site while only a HIS residue forms the M_en_ site. HIS635 forms the M_en_ site while GLU636 forms the M1 site, while GLU640 and ASP643 form the M2 site. In ZIP5, the M1 site contains HIS423 while HIS427 and GLU428 form the M2 site. In ZIP7, GLU359 forms the M1 site. And the second HIS residue HIS362 and GLU363, ASP366 form the M2 site. In ZIP13, the first GLU residue of this motif (GLU258) forms the M1 site while the second GLU residue, GLU262, forms the M2 site (Table 4).

In the predictive models discussed in this article, except ZIP1,2,3, and 9, all other members have a binuclear site. ZIP8 and 14 possibly have a trinuclear site. Besides the zinc ion binding sites in the TMD (M1,2,3 sites), another possible binding site was also found (Table 4). The M_en_ site is located on the top of the TMD. It may help the zinc ion move to the TMD from the ECD. Another potential site is the M_ex_ site which is located at the bottom of the TMD. From the M_ex_ site, the zinc ion would move into the ICD. For the ZIPI, ZIPII, gufA subfamilies, all the members have M_ex_ sites.

#### 3.5.1 ZIP1, 2, 3, 9, and 11

Within this group, except ZIP9, all of them have the GLU-rich area on the top of the TMD, which may help attracting zinc ions in the ECD, and all of them have M_ex_ sites. ZIP11 has a binuclear site in the TMD.

In ZIP1, the zinc ion may be attracted by GLU residues on the long N-terminus containing GLU6,8,15,20,98,209, and 262 in the ECD. At the top of TM2, HIS104 is located. Facilitated by interactions involving the N-terminus and TM2, the zinc ion would be guided to the M_en_ site of the TMD, which comprises ASP91, GLU276, and HIS190. From there, with the assistance of HIS190, the zinc ion proceeds to the M1 site, defined by ASP87 on TM2 and HIS190, GLU194 on TM4, and HIS217 on TM5. Finally, the zinc ion translocation may be driven by the coordinated action of three GLU residues at the base of TM7 (GLU290, 295, and 300) to the M_ex_ site of the TMD, which consists of SER70, GLU295, and CYS74. With the help of GLU residues, the zinc ion may bind with the motif of HWH between TM3 and 4, leaving the TMD and entering the cytoplasm (Figure 9).

**Figure 9.**
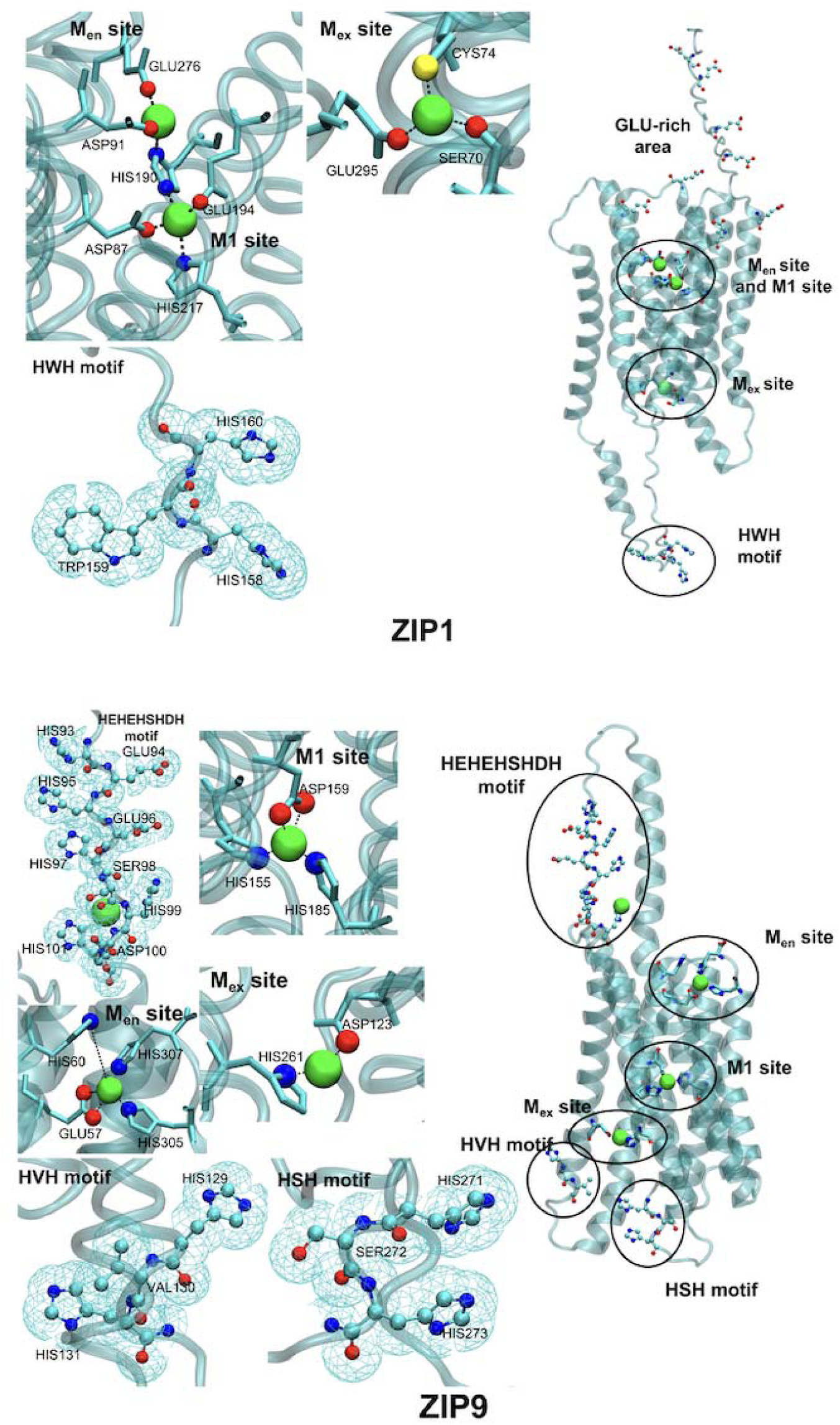
Proposed zinc transport pathway for ZIP1 and 9. Neither of these transporters contain the binuclear metal sites. ZIP1 has a GLU residue rich area to attract zinc ions. The zinc ion will be transported to the M_en_ site composed by GLU276, ASP91, and HIS190 and then reach the M1 site around ASP87, GLU94, HIS127 and 190. There are 3 GLU residues assisting the zinc ion to move out of the TMD. ZIP9 has an HEHEHSHDH motif in the ECD, which may help attracting zinc ions. ZIP9 has only the M1 site composed by HIS155, ASP159, and HIS185.

In ZIP2, the zinc ion may be initially attracted by abundant GLU residues at the top of TM2 and 3, including GLU70 and 73 on TM2 and GLU106 on TM3. These residues may attract the zinc ion from outside the membrane toward a GLU-rich region between TM2 and 7, facilitating its entry into the TMD. The entry site is formed by three GLU residues: GLU262 on TM7 and GLU67,71 on TM2. Once inside the channel, the zinc ion is directed to the M1 site comprising HIS63 on TM2, HIS175, GLU179 on TM4, and HIS202 on TM5. From this site, the zinc ion may be guided by GLU276 and 281 on TM7, HIS40 at the bottom of TM2, and HIS216 at the base of TM5 toward the TMD exit. Finally, the zinc ion may bind with the HSHGH motif on the loop between TM3 and 4, enabling its exit from the TMD (S, Figure 2).

In ZIP3, the zinc ion may initially be captured by a GLU-rich network at the top of the TMD, consisting of GLU192 and 193 on the loop between TM4 and 5, GLU193 at the top of TM5, and GLU251 on TM6. The zinc ion may be first captured by the M_en_ site containing GLU69 and 193. The zinc ion then may be guided to the M1 site with the assistance of GLU184 on TM4 and coordinated by HIS180 and GLU184 on TM4 as well as HIS207 and GLU208 on TM5. From there, it appears that there are two pathways. The zinc ion could bind to GLU280 on TM7 and move to the gate of the channel formed by TM2,7,8, and 3. Alternatively, the zinc ion could move to the M_ex_ site, which is composed of GLU285 and 288. In the intracellular region, HIS33, GLU35 and ASP38 at the end of TM2 may help the zinc ion leaving from the M_ex_ site. The zinc ion may also bind with the HGH motif on the loop between TM3 and 4, allowing it to leave and enter the deep membrane region (S, Figure 3).

In ZIP9, the long loop between TM2 and 3 contains an HEHEHSHDH motif that may help attracting the zinc ion from the ECD, especially HIS99, ASP100, and HIS101. Additionally, the extended TM2 features HIS71 and 72 in the ECD and may also assist in capturing the zinc ion. The zinc ion will move to the M_en_ site, which is composed of GLU57 and HIS60 on the TM2, and HIS305 and 307 at the top of TM8. Once inside the channel, the zinc ion reaches the M1 site coordinated by HIS155 and ASP159 on TM4 and HIS185 on TM5. Guided by HIS261 on TM7, the zinc ion moves from the M1 site to the M_ex_ site within the TMD, which is composed of ASP123 on TM3 and HIS261 on TM7. Finally, the zinc ion may bind to the M_ex_ site with the assistance of the HSH motif on the TM7–TM8 loop and the HVH motif at the base of TM3 (Figure 9).

In ZIP11, zinc ion transport may be primarily facilitated by GLU residues due to the relative scarcity of ASP and HIS residues in this protein. The zinc ion may be first captured by a network of GLU residues in the ECD, including GLU65 on TM2, GLU220 on TM4, GLU228 on TM5, and GLU288 on TM7. ZIP11 has a binuclear site. The M1 site contains HIS204, GLU208 on TM4, GLN240 and GLU244 on TM5. There is also an M2 site composed of ASN205 and GLU208 (on TM4), ASN241 and GLU224 (on TM5). GLU208 is the bridge residue between M1 and M2 sites. The M_ex_ site is composed of ASP96 on TM3, ASP308 and ASP309 on TM7. Finally, possibly with the assistance of HIS101, GLU106, and ASP107 at the base of TMD, the zinc ion connects with residues near the extra α-helix tips between TM3 and 4 in the ICD, exiting the TMD and entering the cytoplasm (S, Figure 4).

#### 3.5.2 ZIP4, 12, 8, and 14

Al of these ZIP transporters have a multinuclear transport site and an ECD. The surface of the ECD contains HIS, GLU and ASP residues that may attract the zinc ion. ZIP8 and 14 potentially have a trinuclear transport site between TM4 and 5. In ZIP8 and 14, there is another potential zinc ion binding site in the transport channel (between TM4, 5 and 7). Also, in the ICD, all four transporters have the HIS motifs that may assist the zinc ion to move into the ICD.

In ZIP4, the zinc ion may be guided by the HSHRH motif on the loop between the third and fourth α-helix of ECD2, as well as the HTH motif between TM2 and 3 of the TMD, which directs it to the TMD entrance. With the help of ASP643 and 644 on TM8, the zinc ion possibly contacts HIS379 on TM2 and then moves further into the channel. The M1 site is composed of ASP375 on TM2, HIS507 on TM4, and HIS536 on TM5. The M2 site is composed of ASP504 on TM4, as well as HIS540, 536, and ASP544 on TM5. Subsequently, the zinc ion may reach the potential M_ex_ site at the base of the TMD, formed by HIS358 on TM2, HIS550 on TM5, and ASP614 on TM8. Finally, the zinc ion may be attracted by the DPEDLED or HSSHSHGGHSH motif on the loop between TM3 and 4 before it exits the protein (Figure 10).

**Figure 10.**
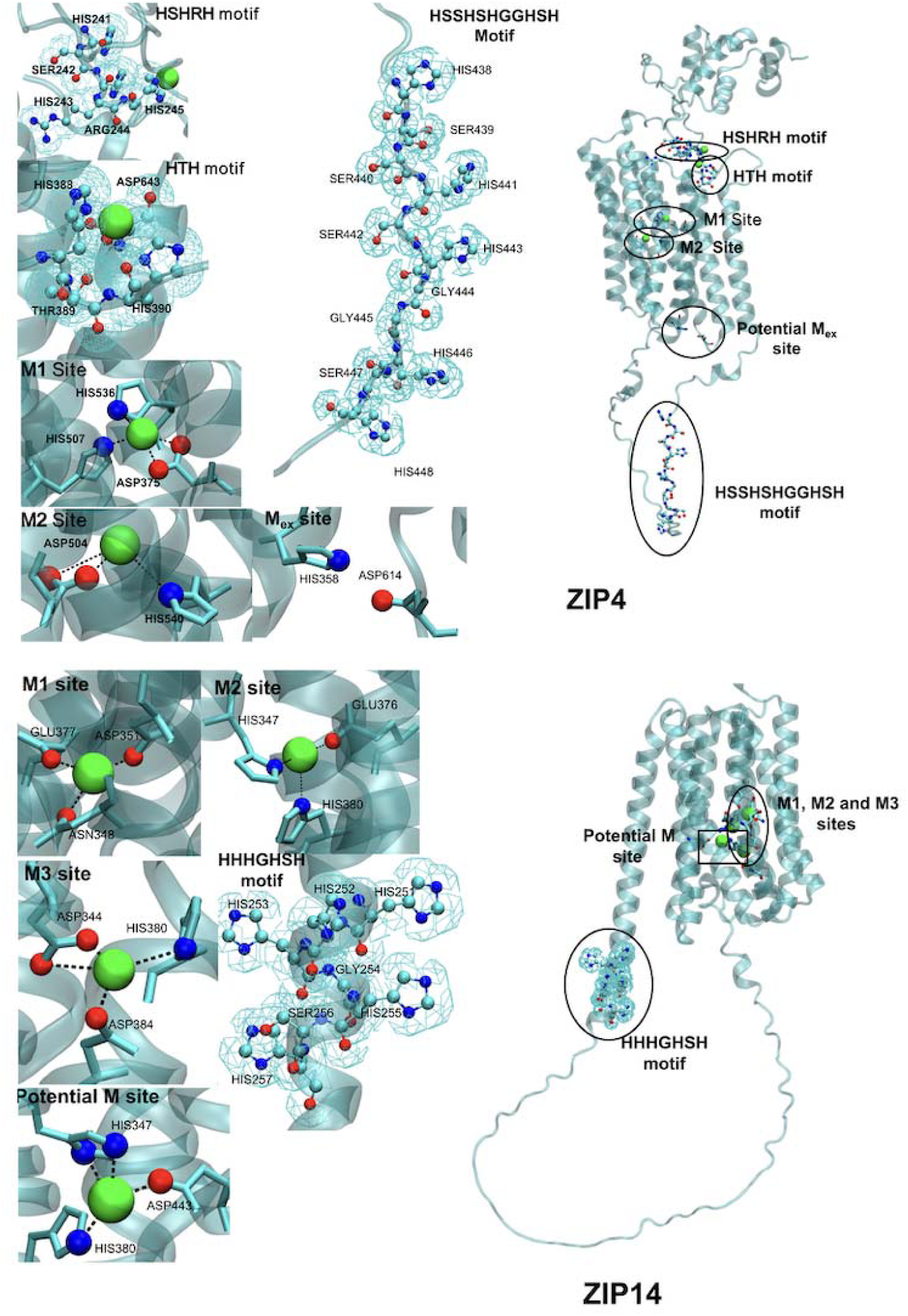
Proposed zinc transport pathway for ZIP4 and 14. For ZIP4, many HIS residues on the ECD may help attracting the zinc ion. First, the zinc ion may bind to the HSHRH and HTH motifs. ZIP4 has two M sites. M1 contains HIS536, 507, and ASP375. M2 contains ASP504 and HIS540. From the M site the zinc ion will move to the M_ex_ site composed of HIS358, 550 and ASP614. Then, with the potential help of the HSSHSHGGHSH motif, which is located between TM3 and 4, the zinc ion will leave the TMD to the ICD. ZIP14 has three M sites. The zinc ion may bind with the HFP motif, which is in the ECD. The M_en_ site is composed of HIS347, GLU376 and HIS380. The M1 site is formed by ASP344 on TM4 and ASP384 on TM5. The M2 is composed of HIS347, GLU376 and HIS380. M3 has ASP344 and ASP384. The zinc ion may be further binding to the HSHH motif located on the loop between TM3 and 4. From there, the zinc ion may be transported to the ICD.

In ZIP12, zinc ions may be attracted by ASP and GLU residues in the ECD. The HFP motif on the loop between TM2 and 3 helps to bind the zinc ion and may direct it to the TMD M_en_ site, formed by HIS428 on the TM2–TM3 loop, HIS419 on TM2, and GLU686 on TM8. M1 has the ligands HIS551 and ASP555 on TM4, and HIS580 on TM5. The ligands at the M2 site include ASP548 on TM4 and HIS584, ASP588 on TM5. GLU648 may facilitate the zinc ion transport from the M1/M2 site. Finally, the zinc ion may bind to the HSHH motif located between TM3 and 4, enabling its transition into the cytoplasm (S, Figure 5).

In ZIP8, a potential zinc ion transport pathway involves initial binding to HIS, ASP, and GLU residues in the ECD. The zinc ion is then guided to a potential M_en_ site on the top of the TMD, formed by ASP189 and 193 on the loop between TM2 and 3. ZIP8 has a trinuclear site. The M1 site is composed of GLU343, 344 and HIS347, which are on TM5, while the M2 site comprises ASP311, ASN315 on TM4, and GLU344 on TM5. The M3 site has ASP311 (on TM4), and HIS347 and ASP351 (on TM5). In the TMD, there is a potential zinc ion binding site between TM4 and 7, which contains CYS310 and HIS314 (on TM4), and ASP410 (on TM7). Finally, the zinc ion may be attracted by the HTH motif on the loop between TM3 and 4, enabling its exit (S, Figure 6).

In ZIP14, GLU, HIS, and ASP in the ECD participate in attracting the zinc ion, which then may bind at the M_en_ site with the assistance of GLU residues clustered on the fifth helical tail of the ECD. ZIP14 has a trinuclear site. The M1 site is composed of ASN348 and ASP351 (on TM4), and GLU377 (on TM5). The M2 site has HIS347 (on TM4), and GLU376 and HIS380 (on TM5). The M3 site contains ASP344 (on TM4), and HIS380 and ASP384 (on TM5). There is another potential zinc ion binding site in the transport channel which contains HIS347 (on TM4), HIS380 (on TM5), and ASP443 (on TM7). After binding to the M sites, the zinc ion may reach the HHHGHSH motif, composed of five HIS residues clustered at the base of TM3, allowing it to exit the TMD and enter the cytoplasm. The elongated TM3 in ZIP14 compensates for the absence of HIS clusters in the TM3–TM4 loop, unlike ZIP8, and instead relies on the HIS-rich motif at the base of TM3 (Figure 10).

ZIP8 and 14 also transport other metal ions. Both transporters contain a conserved EEXPHEXGD motif in the TM5. This sequence, featuring a GLU residue instead of the usual HIS residue, is thought to affect how these transporters recognize and move different metal ions, not just zinc. As a result, ZIP8 and 14 can transport other metals like iron (Fe²⁺), manganese (Mn²⁺), and cadmium (Cd²⁺). In addition, both proteins have HIS-rich segments located in the intracellular loop between TM3 and 4. ZIP14 contains four histidine residues (HX)₄ in this region, while ZIP8 has two (HX)₂. Though the exact role of these HIS clusters is still unclear, they could be involved in metal binding or possibly regulating transporter function. These structural features together help explaining how ZIP8 and 14 can interact with a variety of metal ions.

#### 3.5.3 ZIP6 and 10

ZIP6 and 10 can combine to form a heteromeric structure [55]. But we focus on the homodimer only here. ZIP6 and 10 have the most complicated structures among the family. In the ECD, both structures have the long loop between the α-helices in the ECD which contains HIS, GLU, and ASP motifs. In our predicted models, both structures have the binuclear transport sites. In the ICD, especially on the loop between TM3 and 4, there are lots of HIS motifs, which may be important for the transport specificity of ZIP6 and 10.

In ZIP6, the zinc ion may be attracted by HIS and GLU residues on the ECD and directed toward the TMD through a motif rich in HIS residues on the loop between the second and third α-helix of the ECD, leading to the first M_en_ site with the help of the HSHASHHSHSH motif between TM2 and 3, which is composed of HIS380, 385, and GLU746. From there, with the help of HIS380 and 385, the zinc ion may move to the next M_en_ site formed by ASP372, HIS376 (both on TM2), and HIS635 (on TM5). Potentially guided further by HIS606 and ASP610 (on TM4), the zinc ion reaches the M1 site, comprising HIS606, ASN607, and ASP610 (on TM4), and GLU636 (on TM5). ZIP6 has an M2 site, which contains ASP603 (TM4), and GLU640 and ASP643 (both on TM5). Subsequently, the zinc ion moves toward the TMD’s base and may interact with ASP713, 716, and HIS717 on TM8. Finally, the HHHHDYHH motif on the loop between TM3 and 4 may facilitate the movement to the intracellular region which contains 5 HIS motifs, such as the HHHHH motif (Figure 11).

**Figure 11.**
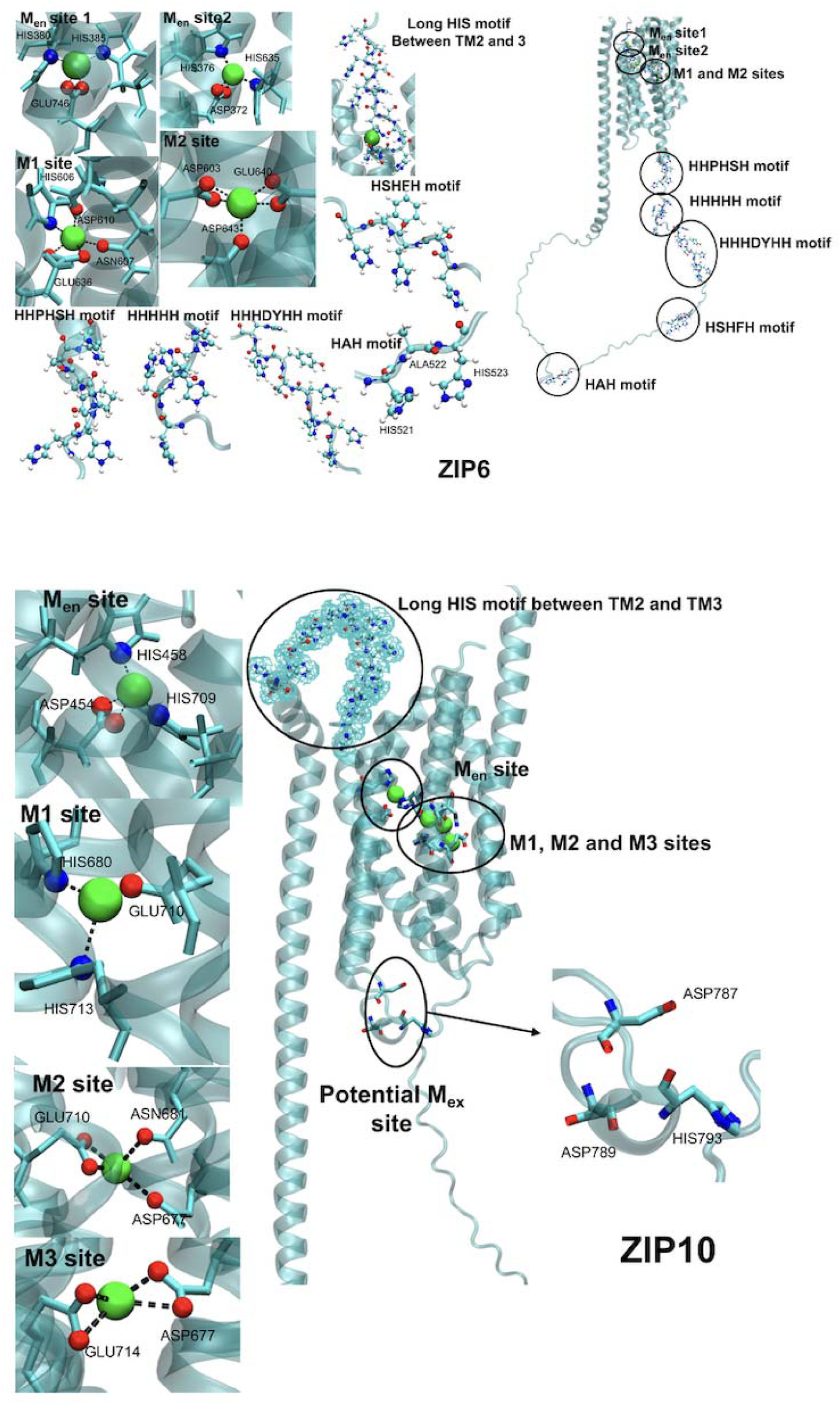
Proposed zinc transport pathway for ZIP6 and 10. Both transporters have a long loop in the ECD which contains many HIS motifs. For these two structures, there is also a long HIS motif on the loop between TM2 and 3. In ZIP6, M1 is composed of ASP603, HIS639, and ASP643. The potential M_ex_ site contains ASP716 and HIS717. In ZIP6, there are 5 HIS motifs on the loop between TM3 and 4. Some of the HIS motifs close to the TMD may assist the zinc ions to move to the ICD. In ZIP10, the M_en_ site contains two HIS residues and one ASP residue (HIS458, 709, and ASP454). In this predicted pathway, ZIP10 has only one M site, which is composed of HIS680 and 713. At the potential M_ex_ site, there are two ASP and one HIS residue (ASP787, 789, and HIS793). There is a long HIS motif on the loop between TM3 and 4 in the ICD.

In ZIP10, there is a long loop in the ECD containing 6 HIS motifs, 3 ASP motifs, and 4 GLU motifs. The zinc ion may bind first to the HDHSHQAHGHSHGH motif between TM2 and 3 before reaching the TMD M_en_ site formed by HIS458 on TM2, and ASP454 and HIS709 on TM5. HIS709 may facilitate the movement to the M1 site, composed of HIS680 on TM4, GLU710 and HIS713 on TM5. ZIP10 also has a trinuclear site. The M2 site has ASP677 and ASN681 (on TM4), and GLU710 (on TM5). The M3 site contains ASP677 on TM4 and GLU714 on TM5. Subsequently, probably aided by ASP787, 789, and HIS793 on the loop between TM7 and 8, the zinc ion could be attracted by the HIS motifs on the loop between TM3 and 4, then exiting the TMD and entering the intracellular space (Figure 11).

#### 3.5.4 ZIP5, 7, and 13

All three have a binuclear metal binding site. In this group, only ZIP5 has an ECD. HIS and GLU residues distributed on the surface of the ECD can attract the zinc ion.

In ZIP5, the zinc ion comes to the M_en_ site formed by HIS264 and 268 on the top of TM2, and GLU530 on the top of TM8. The zinc ion will then move to the M1 site, which consists of HIS394 and ASP398 on TM4 and HIS423 on TM5. There is also the M2 site between TM4 and 5 comprising ASP391 on TM4, GLU428 on TM5. There are two potential binding residues, ASN395 (on TM4) and HIS427 (on TM5), around the M2 site with distances to the zinc ion between 3 and 4 Å. Finally, the zinc ion may reach the HSHGH motif on the loop between TM3 and 4 (Figure 12).

**Figure 12.**
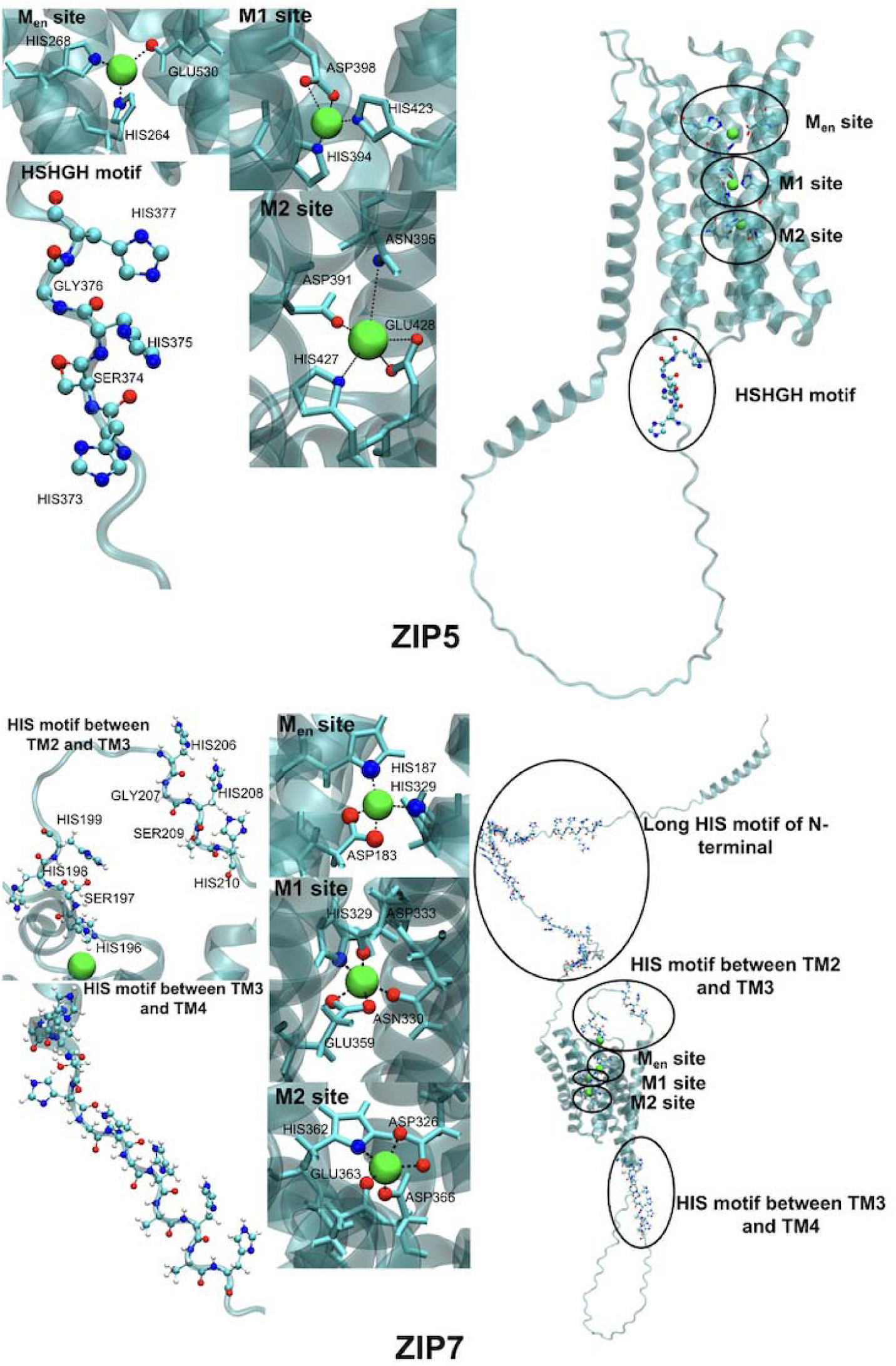
Proposed zinc transport pathway for ZIP5 and 7. ZIP5 has one ECD, which may help attracting zinc ions for transport. The M_en_ site for the zinc ion is composed of HIS264, 268, and GLU530. ZIP5 only has one M site, which is composed of HIS394, ASP398, and HIS423. Then, the zinc ion will be transported to the M_ex_ site, which has ASP391, ASN395, HIS427, and GLU428. There is a HSHGH motif on the loop between TM3 and 4 in the ICD, close to the TMD, possibly moving the zinc ion out of the TMD. ZIP7 has a long N-terminal, which contains many HIS residues that could interact with the zinc ion in the ECD. ZIP7 has two M sites. M1 has HIS329, ASP333, ASN330, and GLU359, while M2 has ASP326, HIS362, GLU363, and ASP366 as ligands. The long HIS motif between TM3 and 4 may assist the zinc ion to move out of the TMD.

In ZIP7, the 5 HIS motifs at the N-terminus may attract the zinc ion. Then, the HGHSH motif on the loop between TM2 and 3 may help the zinc ion to approach the TMD, and a HSHH motif on this loop possibly assists the zinc ion entering the TMD. The zinc ion will bind to HIS196 on TM2 and then move to the M_en_ site composed of HIS187 on TM2, HIS329 on TM4, and HIS358 on TM5. With the help of HIS329 on TM4, the zinc ion will move to the M1 site. The M1 site consists of HIS329, ASP333, and ASN330 on TM4, along with GLU359 on TM5, while the M2 has ASP326 on TM4, and HIS362, GLU363, and ASP366 on TM5. After exiting from the M site(s), HIS residues at the bottom of TM3 may bind the zinc ion with the HGHSHGHAHSH motif before it leaves the protein (Figure 12).

In ZIP13, which is located in an intracellular membrane, a potential zinc ion transport pathway begins with GLU residues at the top of the TMD. GLU128 on the top of TM2 and ASP371 on the top of TM8 assist the zinc ion reaching the TMD entrance. There is a potential M site with one zinc ion bound in the TMD which is composed of ASN120 (on TM2), ASP228 (on TM4), and GLU258 and HIS261 (on TM5). The M1 sites comprising ASP228, ASN229, and HIS232 on TM4, along with GLU258 on TM5 while the M2 site comprises ASN225 on TM4, GLU262 on TM5, and ASP265 and GLN283 on TM6. With further potential support from ASP265 and GLU262 on TM5, the zinc ion may migrate to ASP residues located at the base of TM7 and 8, ultimately exiting the TMD. Additionally, ASP184 and HIS196 on the loops between TM3 and 4 may facilitate the zinc ion’s progression to the ICD (S, Figure 7).

## 4. Discussion

### 4.1 ZIP structures

Human ZIP1 and 2 were cloned 25 years ago, the four subfamilies identified, and the membrane topology with eight α-helices and potential metal-binding sites in TM4 and 5 predicted [56]. Despite numerous and a rapidly increasing number of publications ever since, there is no experimentally derived 3D structure of any of the 14 mammalian ZIP transporters. Therefore, the prediction of the structures of all 14 members of the human ZIP transporter family with the capabilities of AlphaFold3 as presented here provides a unique and valuable resource with a lot of information for future investigations in many areas of the biochemical and biomedical sciences. There is considerable merit in computational approaches viz-a-viz experimentally determined structures. Parts of the proteins with specific secondary structure have high confidence parameters in structural predictions. Other parts that are disordered have lower confidence parameters, however, but they are usually not revealed in experimentally determined structures either. X-ray diffraction or CryoEM structures leave uncertainties as they are models, too, and they are often determined on truncated forms, reflect conformations frozen at low temperature or are restrained by crystal formation, and the experiments sometimes employ supraphysiological concentrations of metal ions or alternative metal ions [57,58].

A structural model of human ZIP4 has been assembled from the structures of the ECD from a mammalian species and a bacterial orthologue (BpZIP), for which structures of both the cadmium-bound and metal-free protein are available [10,12,19]. While all ZIP family members have the TMDs with eight α-helices, the AlphaFold3 predictions show significant variability in the arrangements of the α-helices, and importantly, there is a remarkably rich structural variety in the domains on each site of the membrane, adding a wealth of information about the structures of ZIP proteins. All human ZIP transporters are modelled as homodimers. However, ZIP6 and 10 are known to form also a heteromer, increasing the structural and functional space even further.

The utilization of over a dozen ZIP transporters exemplifies the highly regulated mechanism of zinc homeostasis at the micronutrient level. The reason for this structural variety is that members of these proteins participate in different processes, not only cellular and subcellular and systemic zinc homeostasis, but also controlling zinc-dependent functions of many cellular processes and generating zinc ion signals for cellular regulation. Furthermore, ZIP proteins interact with a host of other proteins that either modulate ZIP protein functions or the functions of the interacting proteins. Presumably, the lack of the interacting partner is one of the reasons for the disordered structural segments in the ZIP proteins, which may adopt defined secondary structures once they interact. In addition to protein-protein interactions and interactions with small molecules (e.g., androgens for Zip9), the loops contain information for endocytosis, e.g. Leu- and Tyr-based motifs on L2 (between TM3 and 4) and recycling, proteolysis, and translocations in cellular trafficking [59]. Some of the transporters need to reach specific membranes in polarized cells and are localized in membranes of mitochondria, lysosomes, endoplasmic reticulum and Golgi apparatus and the nucleus as is the case for Zip11 [60].

### 4.2 Structures of metal coordination in the selectivity filter and the presence of other metal-binding sites

The structural variability also pertains to metal coordination in the selectivity filter and other metal-binding sites in the protein. The basis for this variability is that the ZIP proteins differ in their selectivity for metal ions, with some of them (ZIP8,14) transporting manganese and iron in addition to zinc, and that different zinc ion concentrations need to be maintained in cellular vs subcellular compartments, which is likely reflected by different metal affinities and possibly metal transport kinetics. Also, we do not know what the chemical structure of the substrate is. It is thought not to be “free” zinc ions as their concentrations are vanishingly low, but rather a zinc complex with a LMW ligand or zinc bound to a protein. The coordination requirements of initial binding and later release of zinc on the other side of the membrane likely differ due to different concentrations of zinc and ligands in the cytosol and cellular organelles. Thus, on both delivery sites, i.e., to and from the ZIP transporter, recognition of additional structures such as metallochaperones or specific zinc-requiring proteins in the cell may be required.

Furthermore, this study highlights not only the M sites but also characterizes the entrance and exit sites. Zip transporters operate via an elevator-type conformational change, allowing zinc translocation across membranes [19,20]. The M1 site is essential for zinc transport, while the M2 site serves an auxiliary role, potentially influencing the primary transport site at M1 [17]. BpZIP releases the metal into the periplasm via a combination of hinge motion and vertical sliding in its transport domain. It also has an M3 site, which has a role in regulation of transport through two mechanisms: acting as a metal reservoir under normal conditions and serving as a sensor to inhibit transport during zinc overload. It stabilizes the inward-facing conformation by slowing metal release from the transport site. Zinc at the M3 site is coordinated by two histidine residues (H149, H151) from IL2 along with D144 (TM3) and E276 (TM7) [22].

### 4.3 The selectivity filter

A key structural feature ensuring substrate specificity is the selectivity filter located within TM4 and 5, where conserved HIS, ASP, and GLU residues coordinate metal binding [17,19]. These residues create a metal-binding pocket that determines ion preference based on size, charge, and coordination geometry, allowing Zn²⁺-specific ZIP transporters, e.g. ZIP4, to favor tetrahedral coordination [61], while Fe ²⁺ -transporting ZIPs, e.g. ZIP14, can accommodate octahedral coordination [49]. The selectivity filter is critical for metal homeostasis, preventing non-specific transport and ensuring that essential metal ions are acquired efficiently [44]. Remarkably, our predictions of metal coordination in the selectivity filter are not completely congruent with predictions made on homology modelling from sequence alignments and aided by the structure of a bacterial orthologue. They include ligands from other α-helices, different coordination numbers, variations in the number of metal ions bound, and assignment of bridging ligands. The predicted binuclear sites for human ZIP4 are HIS507, 536 and 540 (ASP511) for the M1 site and ASP511, GLU537 for the M2 site based on homology modelling [17] while our predictions are ASP375, HIS507 and536 for M1 and ASP504, HIS540 for M2 from the structural model. We do confirm changes of ligands, if the selectivity of the ZIP requires accommodation of metal ions other than zinc. We do not know, however, the functional significance of the additional metal ions at M3 and possibly M4 identified here. The interaction and relationship between the different sites require further validation through molecular dynamics simulations.

### 4.4 Other metal-binding sites

The predictions of human ZIP4 based on homology modelling of known partial structures of orthologues include an intracellular loop IL2 containing an HHH motif and an HHH motif on the extracellular site, both thought to be involved in the entry and exit of zinc ions [12]. In our AlphaFold3 predictions, we identify such entry and exit sites in most members of the family and ligand donor atoms of other amino acids. From this information, we propose pathways of translocation. ZIP transporters are metalloproteins in the sense that they have metal-binding sites in addition to the main transport site. We do not know the functions of these sites but as shown for the inhibitory M3 site in BpZIP [22], some of these sites are likely involved in regulating the transport function and thereby regulating zinc supply in the cell and cellular organelles while others may have additional functions in controlling processes such as the transporter processing and trafficking.

### 4.5 Coordination dynamics

In the elevator mechanism, the transport domain (TM1/4/5/6, HB1) moves vertically against the static scaffold domain (TM2/3/7/8, HB2) to enable alternating access for metal transport [22,62]. Molecular dynamics simulations show that the inward-to-outward-facing transition occurs through a mix of vertical sliding and hinge motion [22]. Other additional ligand-binding sites, such as the entrance (M_en_) and the exit (M_ex_) sites, are generally also associated with TM2 and 7.

Unless there are yet to be functionally characterized permanent metal binding sites in the ZIP transporters, all the sites identified are transient as they are either involved in transport or regulation and they underly conformational dynamics of the proteins. ZIP transporters are categorized as solute carrier proteins (SLC). They work by at least three different types of “mechanisms”. They are carriers and/or channels that are operated by co-transport of ions, activated by phosphorylation [63] or they serve as a G-protein coupled androgen receptor [64]. Another hypothesis is that (some) ZIPs are membrane-bound zinc chaperones, providing zinc for regulation of proteins that bind to them at the side facing the cytosol, e.g., Stat3 [65] and GSK-3B [66].

As with experimental structure determinations a limitation of the structural models is that ZIP proteins undergo conformational changes during their transport activity. A wider conformational space needs to be explored and involves also posttranslational modifications such as phosphorylations, ligand-binding induced conformational changes, and the existence of a several transcripts for each of the transporters.

### 4.6 Implications for additional work

The structures presented here serve as a starting point for many types of investigations. Among those are determination of the effect of multiple mutations in each of the ZIP transporters. Since most of the ZIP transporters are involved in disease processes, the structural information can be used for drug design. The structures also indicate which truncated form of the proteins would be suitable for investigations of 3D structure with experimental methods. Future time-resolved computational investigations will need to address some of the limitations of the static model of the metal transport pathway given here.

## 5. Conclusion

The improvements in the predicted 3D structures of the 14 human ZIP proteins, including their representation as dimers, inclusion of zinc ion binding sites, higher resolution, improve significantly the structural information as there are no experimentally determined 3D structures available for mammalian proteins aside from the ECD of Zip4. These advancements make these models a major resource for future investigations of ZIP family proteins as zinc ion and other metal ion transporters in a plethora of physiological and pathological processes.

The findings reveal intricate zinc ion transport mechanisms within the ZIP protein family, characterized by unique structural arrangements and residue distributions across TMDs, ECDs, and ICDs. Key residues—HIS, ASP, and GLU—play pivotal roles in attracting, coordinating, and translocating the zinc ion through distinct transport pathways tailored to each ZIP family member. The variations in arrangements of α-helices, presence of HIS- and GLU-rich motifs, and additional structural features such as disulfide bonds and extended loops highlight the structural and hence functional diversity within the ZIP family. These elements collectively enhance zinc ion binding efficiency, stabilize protein structures, and guide the zinc ion through sequential transport sites to intracellular regions. The comprehensive understanding of these pathways provides a foundation for further experimental exploration of ZIP protein functionality with implications for cellular zinc homeostasis in particular and cell function in general.

## Acknowledgments

N/A

## Data availability

The dataset in the current study is available from the Xilan Wang upon reasonable request.

## Abbreviations

AF: AlphaFold
ECD: extracellular domain
pTM: predicted TM-score
ipTM: interface pTM-score
HB: helix bundle
HRD: histidine-rich domain
ICD: cytoplasmic domain
IL2: second intracellular loop
M_en_: entrance site
M_ex_: exit site
PCD: PAL-containing domain
TM: transmembrane
TMD: transmembrane domain
ZIP: Zrt,Irt-like protein

