## Supplemental Figures for "Structures and zinc ion transport pathways of the human SLC39A family of metal transporters": Metallomics2025_Wang et al._supplFigures.docx

**Supplementary Figures**

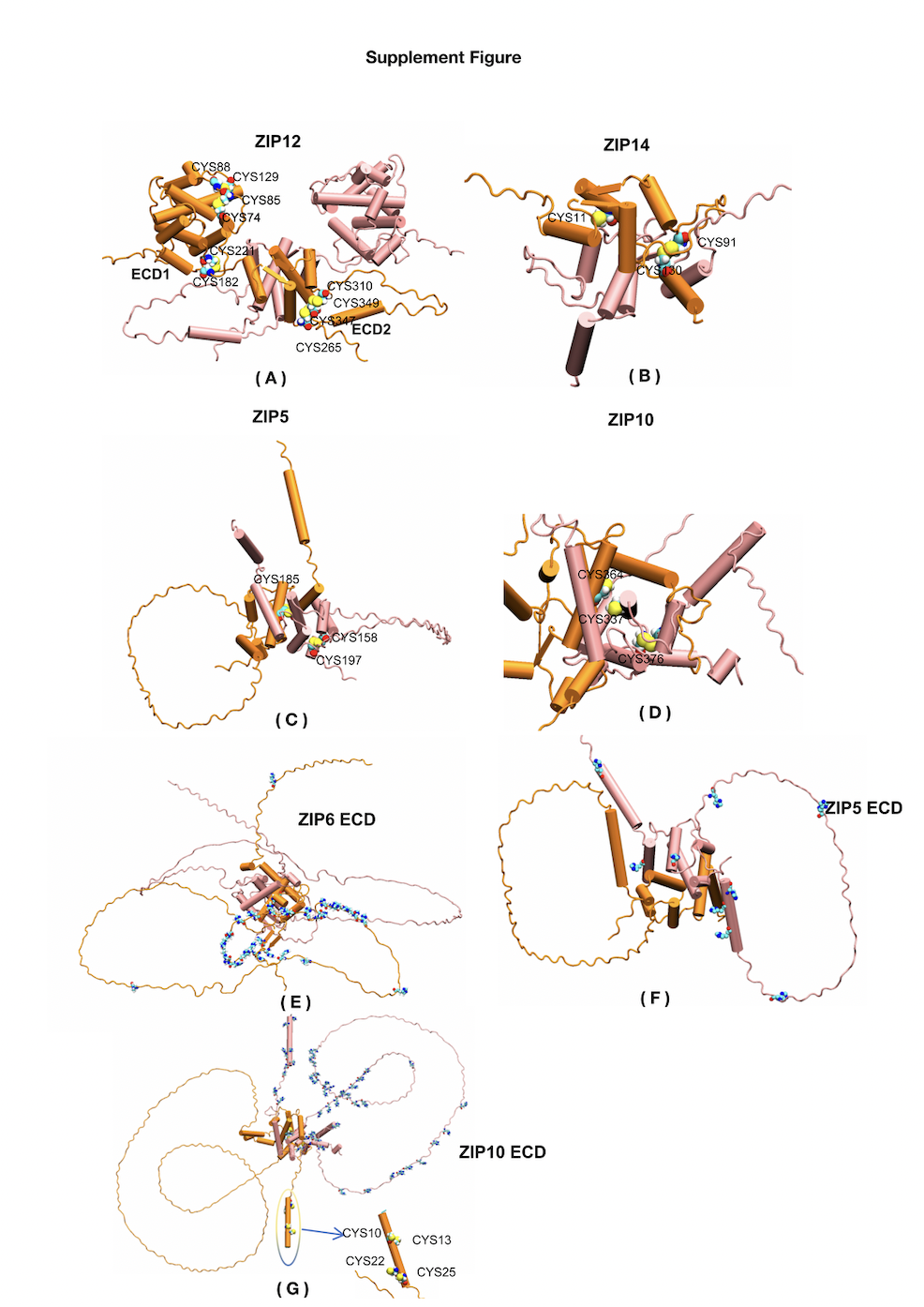

**Supplement Figure 1.** The extracellular domains (ECDs) of ZIP12,14,5,6, and 10, along with the distribution of CYS and HIS residues within the ECDs. The ECD of the first monomer is highlighted in orange, while the ECD of the second monomer is shown in pink. Panels A, B, C, and D display the localization of CYS residues within the ECD monomer of ZIP12,14,5, and 10, respectively. Panels E, F, and G present the histidine (HIS) residue distribution within the ECDs of ZIP6,5, and 10 monomers.

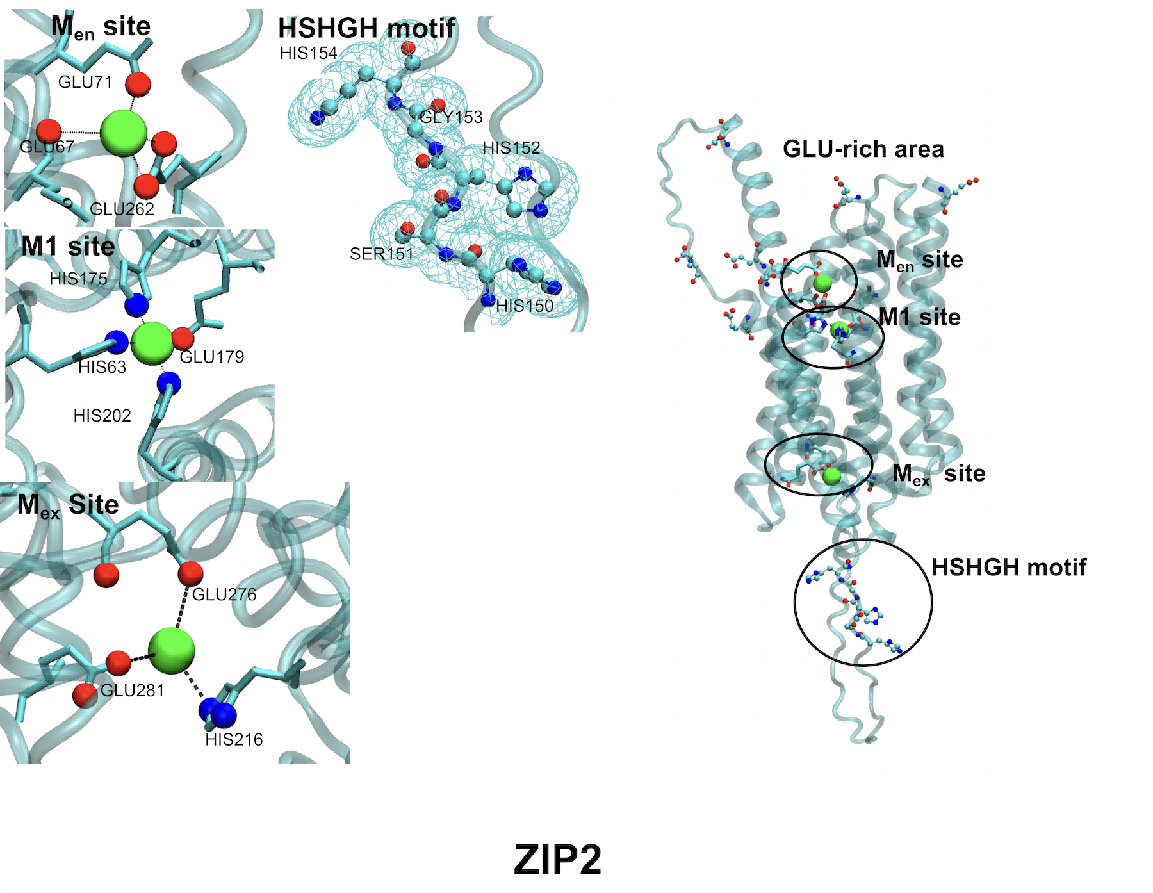

**Supplement Figure 2.** The proposed zinc transport pathway for ZIP2. ZIP2 only has one M site. Zinc ions may be attracted by the GLU residues from the ECD on the top of the TMD. Then the zinc ion will be transported to the M_en_ site composed of GLU71, 67, and 262. The zinc ion will be transported to the M1 site composed of HIS63, 175, GLU179, and HIS202. The M_ex_ site has HIS216, and GLU276, 281. There is also HSHGH motif on the loop between TM3 and 4 to potentially move zinc ions out of the TMD.

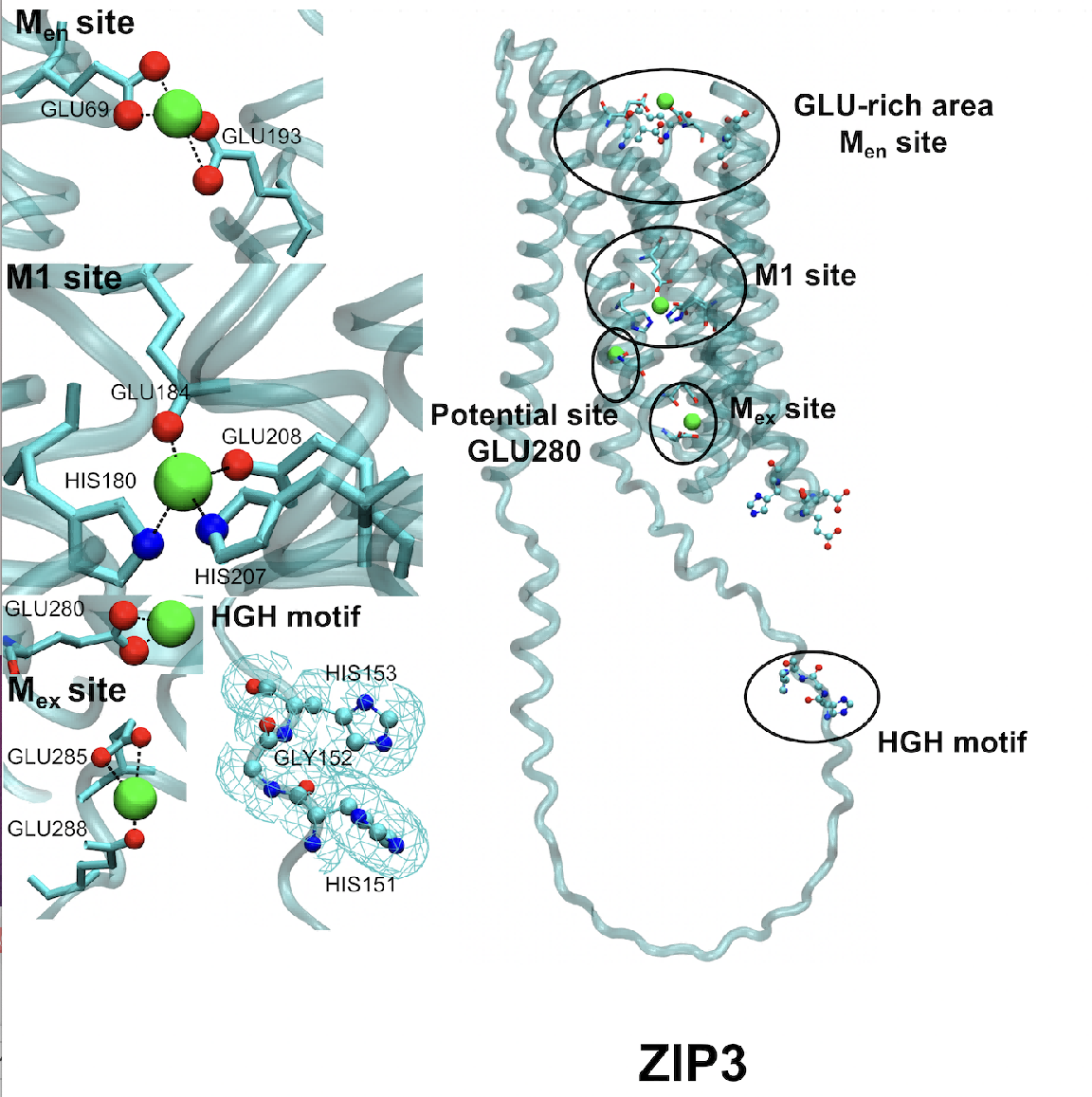

**Supplement Figure 3.** The proposed zinc transport pathway for ZIP3. There are 2 GLU residues to form the M_en_ site (GLU69, 193). After contacting this site, the zinc ion will move to the M1 site composed of GLU184, 208, and HIS180,207. From there, the zinc ion can either move to the GLU280 or to the M_ex_ site composed of GLU285 and 288. There is a HGH motif on the loop between TM3 and 4, which could help the zinc ion to move further to the ICD.

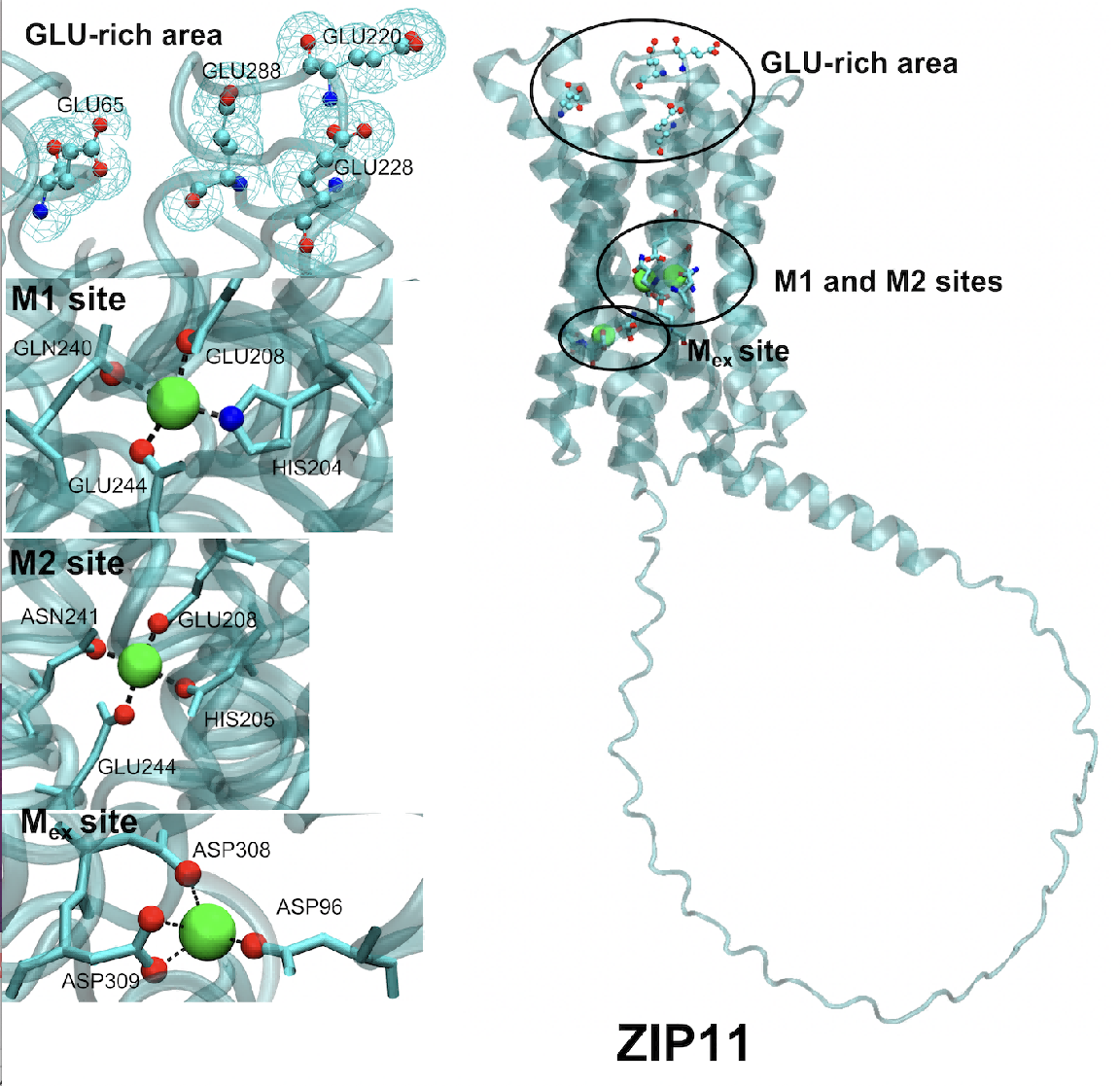

**Supplement Figure 4.** The proposed zinc transport pathway for ZIP11. On the top of the TMD, there are 4 GLU residues to help attracting zinc ions in the ECD. ZIP11 has a binuclear site. The M1 site is composed of HIS204, and GLU208, 244 while the M2 site contains HIS205, GLU208, ASN241, and GLU244. Finally, The M_ex_ site contains three ASP residues: ASP96, 308, and 309. There is no HIS motif in the ICD but the GLU residues will help the zinc ion moving out of the TMD.

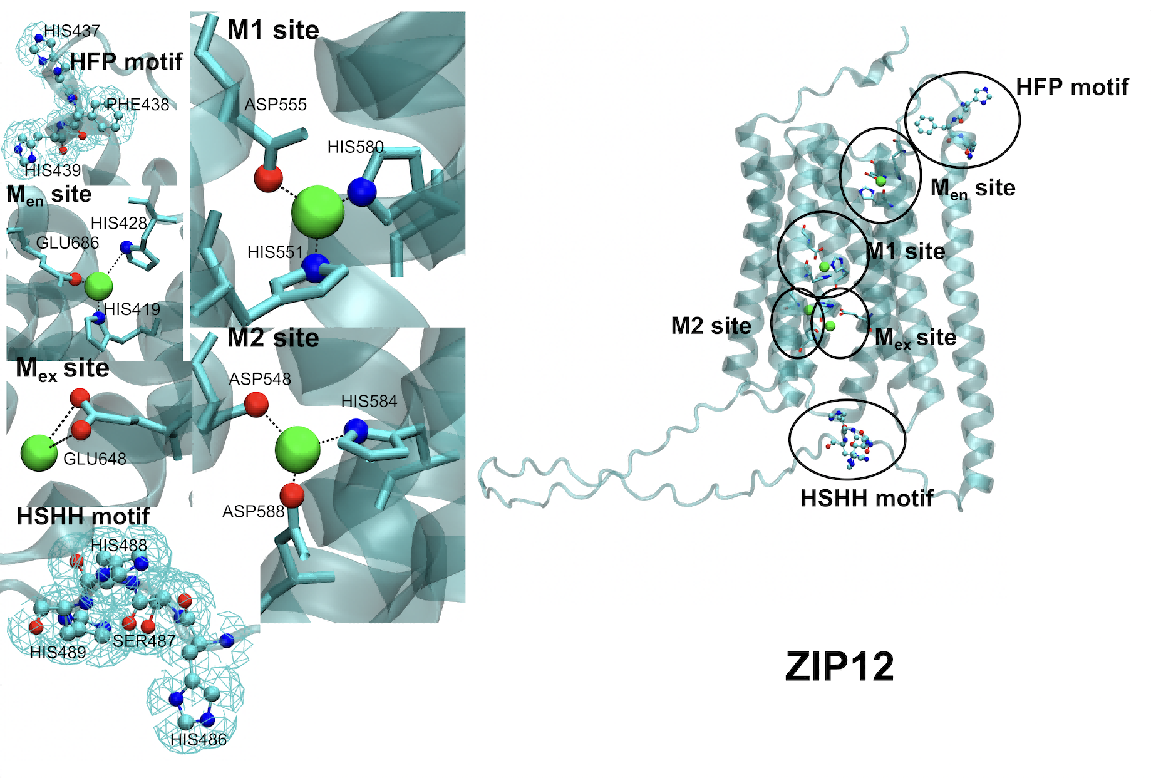

**Supplement Figure 5.** The proposed zinc transport pathway for ZIP12. The ECD may attract zinc ions. The HFP motif may help the zinc ion moving down to the M_en_ site composed of HIS419, 428, and GLU686. ZIP12 has two M sites. The M1 has HIS551, ASP555, and HIS580. The M2 is formed by ASP548, HIS584, and ASP588. After binding to the M site, the zinc ion may be transported to the M_ex_ site formed by GLU648. The HSHH motif may help the zinc ion moving out of the TMD.

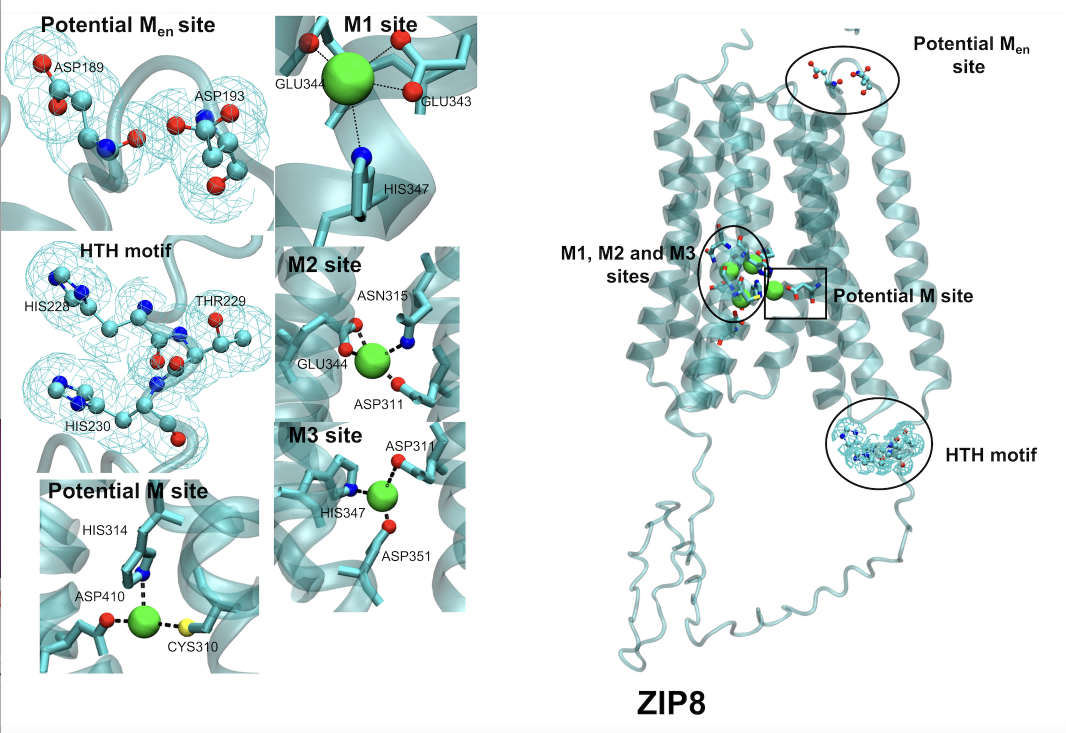

**Supplement Figure 6**. The proposed zinc transport pathway for ZIP8. Two ASP residues may form the M_en_ site (ASP189, 193). ZIP8 has three M sites. The first one is composed of GLU343, 344, and HIS347 and the second one has ASP311, ASN315, and GLU344. The M3 site contains ASP311, HIS347, and ASP351. On the loop between TM3 and 4, the HTH motif may move the zinc ion out of the TMD.

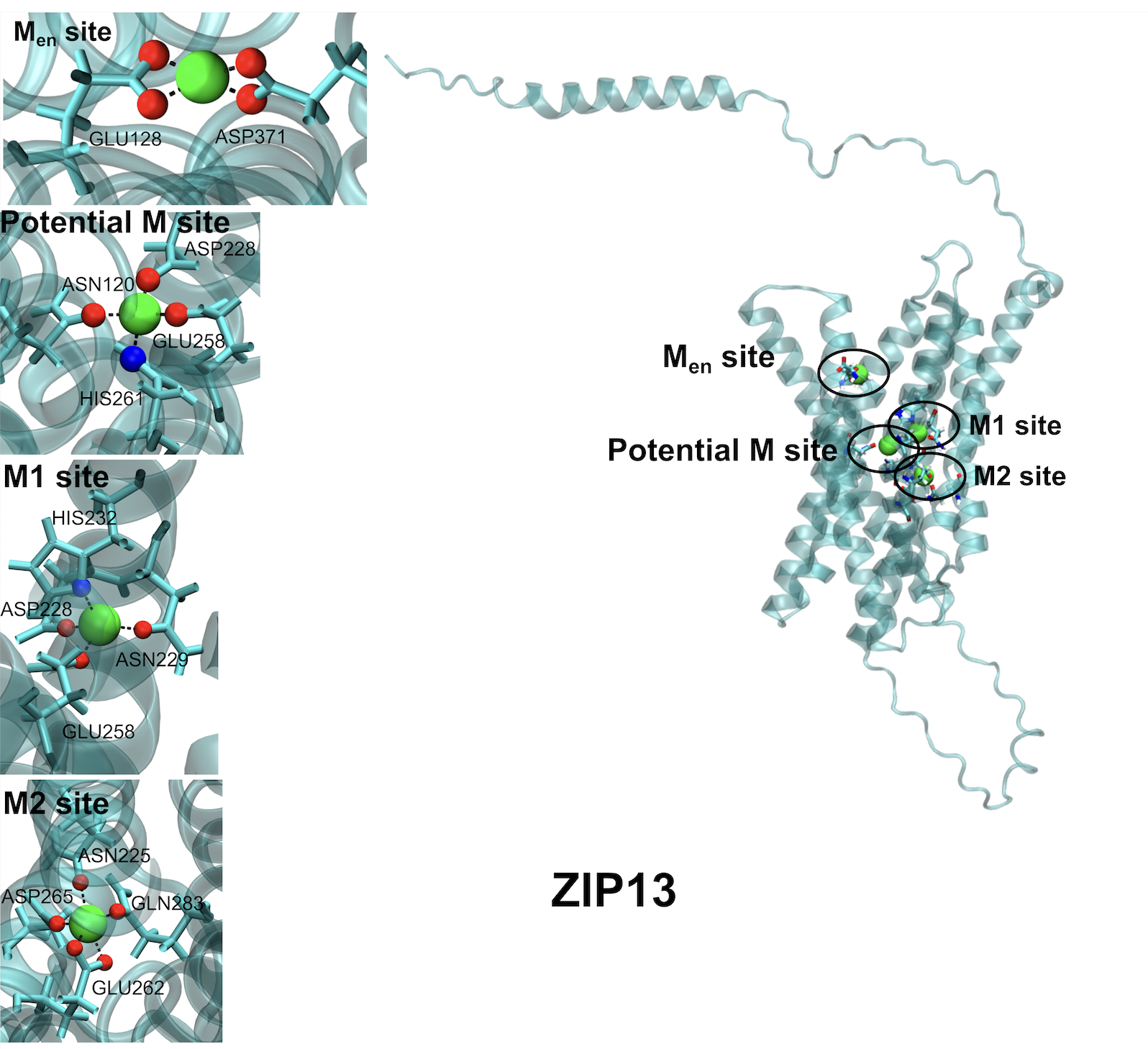

**Supplement Figure 7**. The proposed zinc transport pathway for ZIP13. GLU128 and ASP371 form the M_en_ site. ZIP13 has two M sites. M1 is composed of HIS232, ASP228, GLU258, and ASN229. The M2 site is formed by ASN225, GLU262, ASP265, and GLN283. There is a potential M site in ZIP13, which contains ASN120, ASP228, GLU258, and HIS261. GLU and ASP residues may help the zinc ion moving out of the TMD. There is no HIS motif on the loop between TM3 and 4.
